# Per1/Per2-Igf2 axis-mediated circadian regulation of myogenic differentiation

**DOI:** 10.1101/2020.07.29.209312

**Authors:** Nobuko Katoku-Kikyo, Ellen Paatela, Daniel L. Houtz, Britney Lee, Dane Munson, Xuerui Wang, Mohammed Hussein, Jasmeet Bhatia, Seunghyun Lim, Ce Yuan, Yoko Asakura, Atsushi Asakura, Nobuaki Kikyo

## Abstract

Circadian rhythms regulate cell proliferation and differentiation but circadian control of tissue regeneration remains elusive at the molecular level. Here, we show that proper myoblast differentiation and muscle regeneration are regulated by the circadian master regulators Per1 and Per2. Depletion of Per1 or Per2 suppressed myoblast differentiation *in vitro* and muscle regeneration *in vivo*, demonstrating their non-redundant functions. Both Per1 and Per2 were required for the activation of *Igf2*, an autocrine promoter of myoblast differentiation, accompanied by Per-dependent recruitment of RNA polymerase II, dynamic histone modifications at the *Igf2* promoter and enhancer, and the promoter-enhancer interaction. This circadian epigenetic priming created a preferred time window for initiating myoblast differentiation. Consistently, muscle regeneration was faster if initiated at night when *Per1*, *Per2*, and *Igf2* were highly expressed compared with morning. This study reveals the circadian timing as a significant factor for effective muscle cell differentiation and regeneration.

**eTOC Summary:** Katoku-Kikyo et al. show that the circadian master regulators Per1 and Per2 control the efficiency of myoblast differentiation via Igf2 activation. This pathway creates a preferred circadian time window for myoblast differentiation *in vitro* and muscle regeneration *in vivo*.

## Introduction

Regulation of mammalian circadian rhythms is centered around the Clock/Bmal1 transcription factor complex (Gustafson and Partch, 2015; Hirano et al., 2016; Papazyan et al., 2016; Takahashi, 2017a). The complex binds the E-box (5’-CANNTG-3’) in promoters and enhancers of thousands of genes to activates their transcription, including the *Cry* (*Cry1* and *Cry2*) and *Per* (*Per1*-*Per3*) genes. Gradually accumulated Cry and Per in turn bind Clock/Bmal1 on DNA and repress its transcription activity, forming a negative feedback loop. Subsequent phosphorylation and ubiquitination of Cry and Per lead to their degradation, reactivating Clock/Bmal1. This oscillating activity of Clock/Bmal1 creates transcriptional circadian rhythms in more than 20% of the genes in the genome in at least one tissue in the body. In addition, Clock/Bmal1 activates retinoic acid receptor-related orphan receptor proteins (RORα-RORγ) and reverse orientation c-erb proteins (Rev-erbα and Rev-erbβ), which compete for the retinoic acid-related orphan receptor response element (RORE) in the *Bmal1* promoter. Opposing activities of ROR as an activator and Rev-erb as a repressor of Clock/Bmal1 form the second circadian feedback loop. The feedback loops in the suprachiasmatic nucleus (SCN) in the hypothalamus are entrained by the light signal transmitted from the retina as the primary external cue (zeitgeber). The SCN clock plays a dominant role (central clock) in synchronizing the feedback loops in other tissues (peripheral clocks), including skeletal muscle. In addition, the peripheral clocks are also entrained by various physiological factors such as body temperature, feeding time, and physical activity.

Circadian regulation is tightly integrated into the genetic program of muscle cell differentiation as demonstrated by several studies (Harfmann et al., 2015; Lefta et al., 2011; Mayeuf-Louchart et al., 2015). First, more than 2,000 genes, including the master myogenic regulators *MyoD* and *Myogenin* (*Myog*), show circadian oscillation in abundance (Harfmann et al., 2015; McCarthy et al., 2007; Miller et al., 2007; Pizarro et al., 2013). Second, whereas Bmal1 promotes satellite cell proliferation and differentiation, and is required for muscle regeneration (Chatterjee et al., 2013; Chatterjee et al., 2015), Rev-erbα acts as an inhibitor of these processes (Chatterjee et al., 2019). Third, Clock/Bmal1 binds the E-box in the core enhancer of *MyoD* in a circadian manner; MyoD then binds the *Bmal1* enhancer and increases the amplitude of *Bmal1* expression, forming a feed-forward loop in myogenesis (Andrews et al., 2010; Hodge et al., 2019). Finally, we previously showed that Cry2 promotes myoblast proliferation and fusion during differentiation in a circadian manner through stabilization of mRNAs encoding cyclin D1, a G1/S phase transition driver, and Tmem176b, a transmembrane regulator for myogenic cell fusion (Lowe et al., 2018).

Mouse Per1 and Per2 share 73.4 % sequence similarity at the amino acid level but are not functionally redundant. *Per1^-/-^* and *Per2^-/-^* mice are grossly normal and fertile; however, both knockout (KO) mice exhibit circadian periods up to 2 hr shorter than wild-type (WT) mice and eventually become arrhythmic in constant darkness (Bae et al., 2001; Cermakian et al., 2001; Zheng et al., 2001; Zheng et al., 1999). In contrast, *Per1^-/-^:Per2^-/-^* mice become arrhythmic immediately after transfer to constant darkness although they are morphologically normal and fertile. As for muscle phenotypes, *Per2^-/-^* mice show a 20% shorter running distance with a treadmill test compared with WT and *Per1^-/-^* mice although the length, weight, contractility, and abundance of several contractile proteins in the tibialis anterior (TA) muscle were similar in the three genotypes (Bae et al., 2006). Currently, virtually nothing is known about whether and how *Per* genes contribute to myogenic differentiation and muscle regeneration.

The present study uncovered insulin-like growth factor 2 (Igf2) as a critical link between Per1/Per2 and myoblast differentiation. Igf2 is a necessary and well-characterized autocrine differentiation promoter of myoblasts that increases in secretion levels during differentiation (Duan et al., 2010; Florini et al., 1991; Yoshiko et al., 2002). Igf2 is also upregulated upon muscle injury and enhances regeneration (Keller et al., 1999; Kirk et al., 2003; Levinovitz et al., 1992). Additionally, several single nucleotide polymorphisms of the human *IGF2* genes are associated with a loss of muscle strength following strenuous exercise (Baumert et al., 2016; Devaney et al., 2007). Whole-body *Igf2* null mice display impaired growth at birth but subsequently grow normally (Baker et al., 1993). Igf2 binds the type I Igf1 receptor (Igf1r) with the highest affinity among several receptors, resulting in its auto-phosphorylation and subsequent activation of the PI3K/AKT pathway and the RAS/MAP kinase pathway (Siddle, 2011; Taniguchi et al., 2006). In particular, activation of p38α/β MAPK by phosphorylation is an essential downstream effector for the promotion of myoblast differentiation by Igf2 (Knight and Kothary, 2011; Segales et al., 2016b). p38 achieves the pro-differentiation function by triggering cell cycle exit, activating myogenic transcription factors, and opening the chromatin of muscle gene promoters (see (Gardner et al., 2015; Segales et al., 2016a) for references). Most Igf2 in blood and local tissues is bound by Igf-biding proteins (IGFBP1-IGFBP7), which up- or downregulate Igf2 functions (Allard and Duan, 2018). For example, whereas IGFBP-3 inhibits myoblast differentiation (Huang et al., 2016), IGFBP-5 is induced during early myoblast differentiation and amplifies the auto-regulatory loop of Igf2 expression, resulting in promoted differentiation (Ren et al., 2008). The current work demonstrates the Per1/Per2-Igf2 axis as a novel pathway underlying myogenesis and uncovered the presence of circadian epigenetic priming at the *Igf2* gene during myoblast differentiation.

## Results

### Disrupted muscle regeneration and myoblast differentiation by Per depletion

To study the roles of Per1 and Per2 in skeletal muscle regeneration, TA muscle in *Per* KO mice was injured by barium chloride injection and their regeneration was assessed by a series of histological analyses. The mice were entrained at 12 hr-light (Zeitgeber Time (ZT) 0-ZT12) and 12 hr-dark cycles (ZT12-ZT24) as diurnal rhythms for two weeks before experiments. TA muscle was damaged and harvested at ZT14; ZT14 (20:00) was selected to compare with the result of the antiphase time point (ZT2, 8:00) later. Hematoxylin eosin (HE) staining on day 4.5 post-injury demonstrated smaller myofibers with centrally-located nuclei, an indication of newly formed myofibers, in *Per1^-/-^*, *Per2^-/-^*, and particularly *Per1^-/-^:Per2^-/-^* mice compared with WT mice (Fig. 1, A-C). This trend continued at least until day 14 and was also observed in uninjured myofibers (Fig. S1, A and B). In addition, myofibers expressing embryonic myosin heavy chain (eMHC), a marker for newly generated myofibers, were smaller in *Per2^-/-^* and *Per1^-/-^:Per2^-/-^* mice than those in WT and *Per1^-/-^* mice (Fig. 1, D and E). Furthermore, the kinetics of cell cycle exit of satellite cells in the KO mice were different from those in WT mice. During muscle regeneration, activated satellite cells re-enter the cell cycle [EdU (5-ethynyl-2’-deoxyuridine)(+)/MyoD(+) population], followed by exit from the cell cycle [EdU(-)/MyoD(+)] before terminal differentiation. Comparison of the frequency of each population on day 4.5 showed an increased frequency of EdU(+)/MyoD(+) cells in the single and double KO mice compared with WT mice (Fig. 1, F and G). This finding could suggest delayed cell cycle exit of the satellite cells in the KO mice, which translates into delayed muscle regeneration on day 4.5. Finally, the single and double KO mice contained more fibrosis on day 14 after the injury as demonstrated by Sirius red stain, suggesting more extensive damage or delayed regeneration in the KO mice (Fig. 1, H and I). Uninjured *Per2^-/-^* and *Per1^-/-^:Per2^-/-^* mice already contained more fibrosis than WT mice although it was less severe than in day 14 mice, implying that natural turnover of myofibers was also disrupted in the KO mice. Thus, both Per1 and Per2 are necessary for the proper regeneration of TA muscle and they can partially compensate each other, a reminiscence of the milder arrhythmic phenotype in the single KO mice than in the double KO mice.

**Figure 1.**
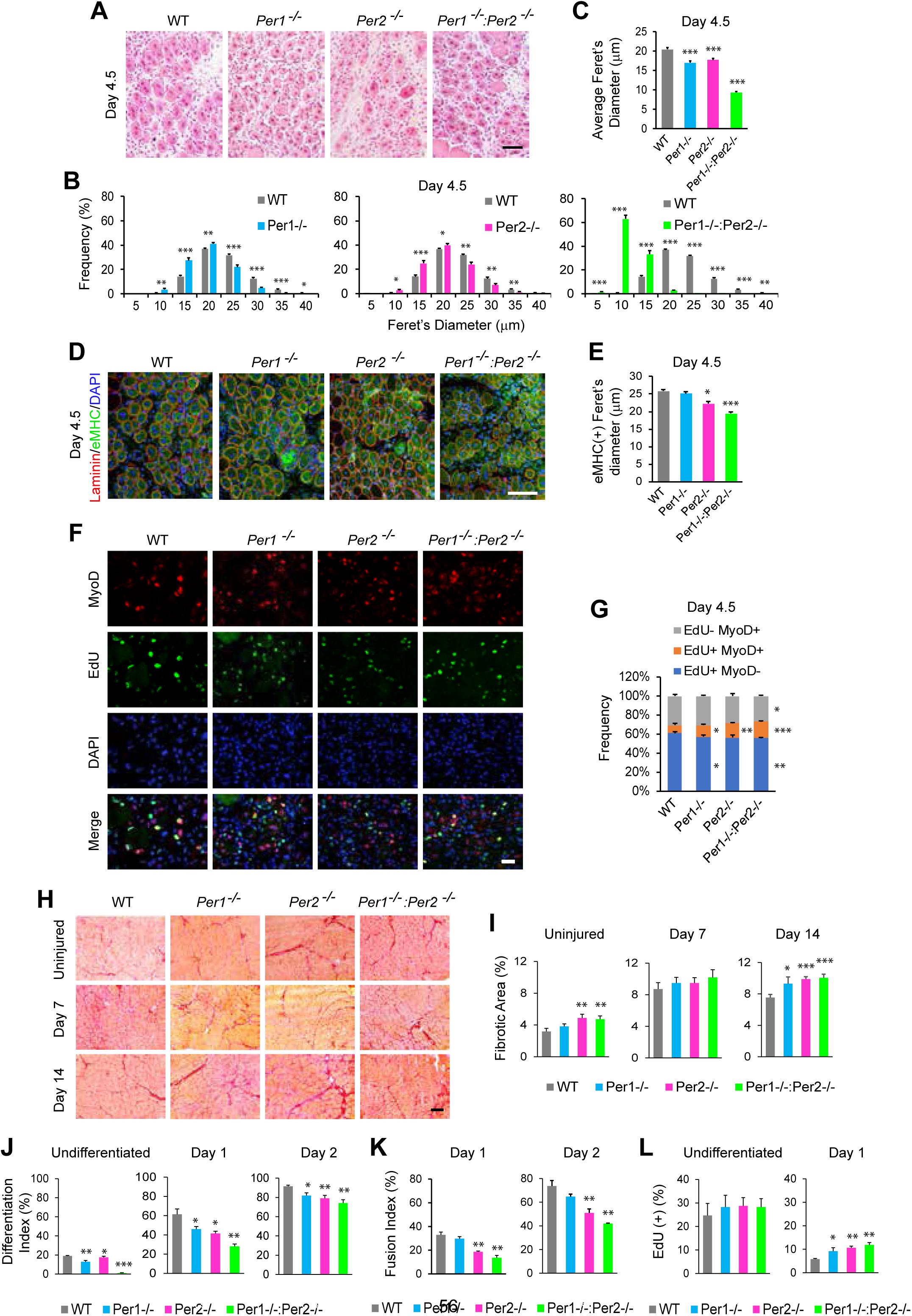
Regeneration of TA muscle in *Per1^-/-^*, *Per2^-/-^*, and *Per1^-/-^:Per2^-/-^* mice. (A) HE staining of day 4.5 TA muscle sections. TA muscle was injured with barium chloride at ZT14 on day 0 and EdU was intraperitoneally injected 96 hr later for (F) and (G). The muscle was harvested 12 hr later as day 4.5. Bar, 100 μm. (B) Size distribution of HE-stained myofibers containing centrally-located nuclei on day 4.5. The minimal Feret’s diameter in each myofiber was measured. n = 8 mice in each group, including 4 males and 4 females, in (B) and (C). (C) Average of the minimal Feret’s diameters of myofibers with centrally-located nuclei on day 4.5. (D) Immunofluorescence staining of TA muscle with antibodies against eMHC and laminin (shows the border of each myofiber) on day 4.5. DNA was counterstained with DAPI. Bar, 100 μm. (E) Average of the minimal Feret’s diameters of eMHC(+) areas on day 4.5. n = 4 mice. (F) Immunofluorescence staining of TA muscle sections with the MyoD antibody and an EdU kit. Bar, 25 μm. (G) Frequency of positive cells for EdU uptake and MyoD staining shown in (F). n = 4 mice. (H) Sirius red staining of days 7 and 14 post-injury and uninjured TA muscle. Bar, 200 μm. (I) The area percentage of Sirius red(+) fibrosis in the groups shown in (H). (J)-(L) Differentiation index (J), fusion index (K), and the frequency of EdU uptake (L) during the differentiation of primary myoblasts prepared from WT and *Per* KO mice. Data represent biological triplicates with n = 80-130 nuclei in each replicate. Data are presented as mean + SEM. * p < 0.05, ** p < 0.01, and *** p < 0.001 with Student’s t-test in comparison to WT.

To understand cell-autonomous effects of *Per* KO, primary myoblasts (activated satellite cells) were purified from hind limbs and induced to differentiate into myotubes with 5% horse serum *in vitro*. *Per1^-/-^*, *Per2^-/-^*, and *Per1^-/-^:Per2^-/-^* myoblasts displayed delayed activation of MHC, a marker for differentiation (Bader et al., 1982). This was quantified as decreased differentiation index (frequency of nuclei in MHC(+) cells among total nuclei) (Fig. 1 J and S1, C-F). These cells also demonstrated decreased fusion index (frequency of nuclei in MHC(+) cells containing more than one nuclei among total nuclei) and increased frequency of EdU uptake compared with WT cells (Fig. 1, K and L, and S1, C-F). These results exhibited impaired cell-autonomous myoblast differentiation by *Per* KO, which could explain the delayed TA muscle regeneration although niche effects could not be excluded.

To obtain a large number of cells for a mechanistic study, we examined whether the mouse myoblast cell line C2C12 could recapitulate the KO phenotypes of the primary myoblasts. The *Per1* or *Per2* gene was depleted by shRNA-mediated knockdown (KD) and CRISPR-Cas9- mediated KO (Fig. S2, A-C). We used KD and KO because we could identify only one effective shRNA clone for each gene. These cells were used in bulk without cloning since differentiation-resistant cells would be selected by cloning. When the cells were induced to differentiate, MHC(+) cells became shorter and more sparse in the KD and KO cells than control cells on differentiation days 3 and 5 (Fig. 2 A). This finding was quantified as decreased differentiation index and fusion index in the KD and KO cells (Fig. 2, B and C). The KO cells also demonstrated slightly delayed cell cycle exit during differentiation (Fig. 2 D). Additionally, expression of differentiation-specific genes encoding *Myog*, muscle creatinine kinase (*Ckm*), myomaker (*Mymk*), and myosin heavy chain-3 (*Myh3*) was decreased by the KD and KO (Fig. 2 E). The similarity of the consequence of the Per 1 and Per 2 depletions was further highlighted by several transcriptome data sets. The list included a heat map, a principal component analysis, Venn diagrams of differentially expressed genes (more than 2,000 genes were commonly up- or downregulated more than 2-fold compared with control cells), and scatter plots (R^2^>0.97) (Fig. 2, F and G, and S2, D-F). In addition, the gene ontology analysis demonstrated that many myogenesis-related genes were commonly downregulated by *Per1* KO and *Per2* KO in undifferentiated cells and differentiating cells on days 3 and 5 (Fig. S2 G), further underscoring the requirement of both *Per1* and *Per2* for proper myogenesis.

**Figure 2.**
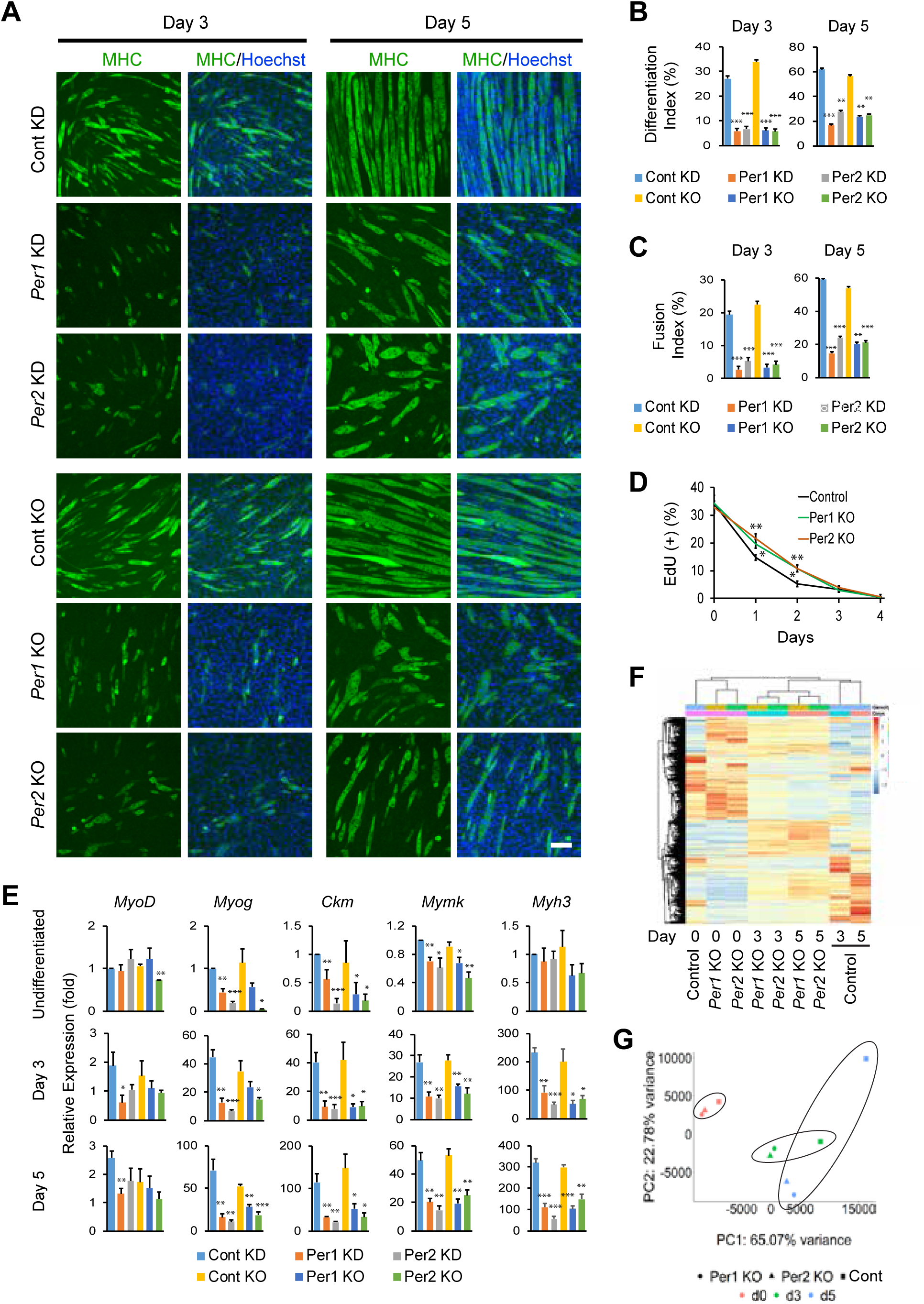
Differentiation of C2C12 cells after depletion of *Per1* and *Per2*. (A) Immunofluorescence staining of C2C12 cells with MHC antibody during differentiation with 5% horse serum. *Per1* and *Per2* were depleted with shRNA (KD) and CRISPR-Cas9 (KO). Non-targeting sequences were used as each control. Bar, 100 μm. (B) and (C) Differentiation index (B) and Fusion index (C) on days 3 and 5. (D) Temporal profile of the frequency of EdU(+) nuclei in KO cells during differentiation. (E) Relative expression levels of five muscle genes determined by qPCR during differentiation. The value obtained with undifferentiated control cells treated with scrambled shRNA was defined as 1.0 for each gene. (F) Heat map comparing the transcriptome of KO cells. Day 0 refers to undifferentiated cells in (F) and (G). n = 1 in (F) and (G). (G) Principal component analysis of KO cells. Data are presented as mean + or ± SEM of biological triplicates unless indicated otherwise. Each replicate includes n = 1,000-1,500 nuclei in (B)-(D). * p < 0.05, ** p < 0.01, and *** p < 0.001 with Student’s t-test in comparison to control.

### Essential role of Igf2 in myoblast differentiation and muscle regeneration

The common phenotypes of *Per1* and *Per2* depletion led us to search for important myogenic genes that were commonly dysregulated by each KO. RNA-seq revealed four muscle genes commonly downregulated in *Per1* KO and *Per2* KO C2C12 cells in an undifferentiated state and during differentiation on days 3 and 5 (Fig. S3 A, highlighted in yellow). They included *Cav3* (encoding the membrane protein caveolin 3), *Csrp3* (the multi-functional protein cysteine and glycine-rich protein 3), *Igf2*, and *Myoz1* (a signaling protein). We decided to focus on *Igf2* because of its significance in myogenesis as described earlier. Downregulation of *Igf2* in *Per* KD and KO cells was verified by qPCR (Fig. S3 B). To investigate the involvement of *Igf2* in the *Per* depletion phenotypes, *Igf2* was knocked down with two shRNAs in C2C12 cells (Fig. S3 C). MHC(+) cells in the KD cells were more sparse and shorter than control KD cells, consistent with the lower differentiation index and fusion index, as well as the decreased expression of differentiation-specific genes (Fig. S3 D, S4 A-C). Cell cycle exit was also delayed by *Igf2* KD during differentiation (Fig. S4 D). Thus, *Igf2* depletion recapitulated the phenotypes of *Per* depletion. Additionally, *Igf2* KD downregulated *Cav3*, *Csrp3*, and *Myoz1*, indicating that these genes were under the control of Igf2 (Fig. 3A).

**Figure 3.**
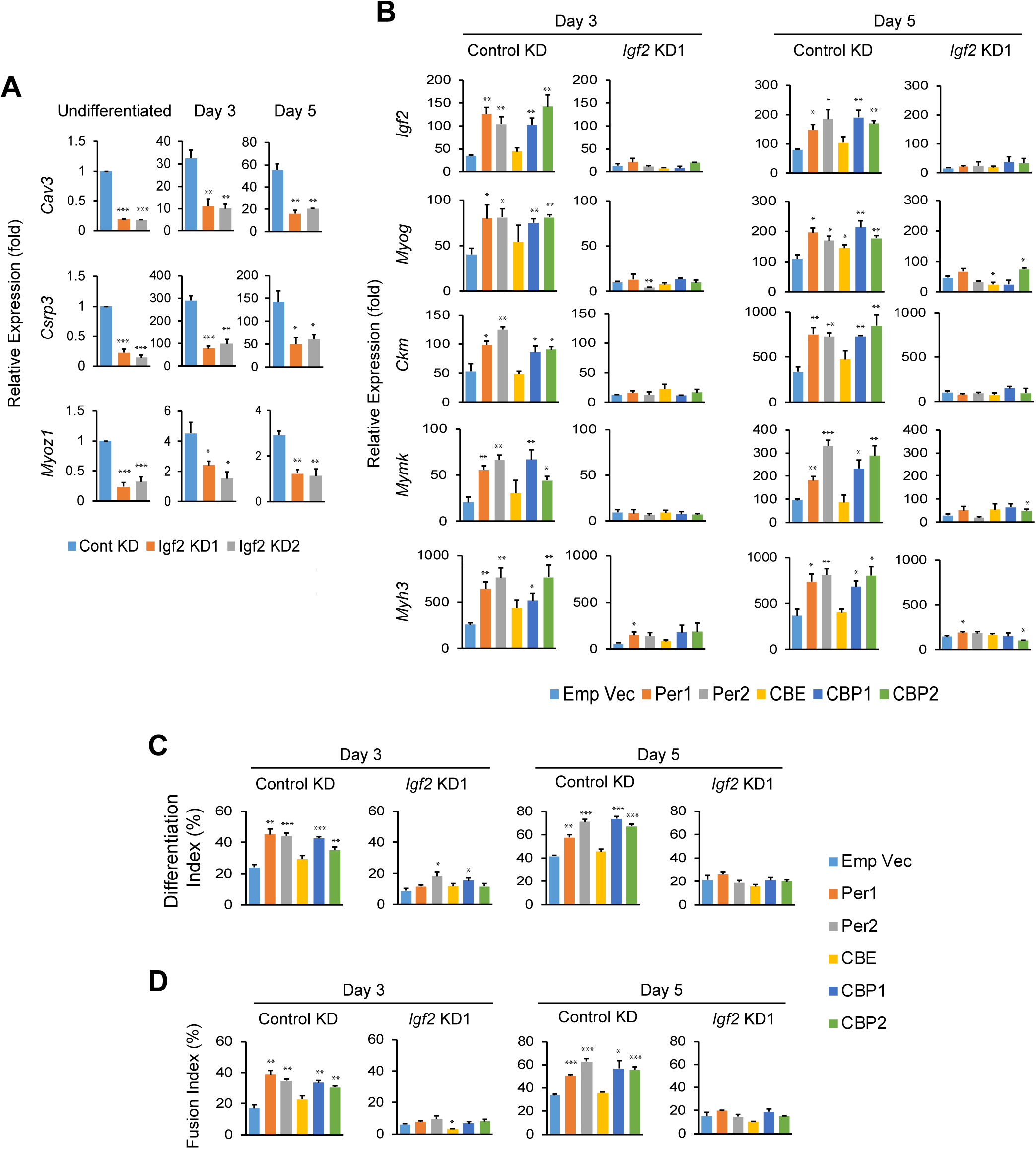
Disrupted differentiation of C2C12 cells by *Igf2* KD. (A) Relative expression levels of *Cav3*, *Csrp3*, and *Myoz1* in *Igf2* KD cells. (B) Relative expression levels of *Igf2* and muscle genes during differentiation of C2C12 cells. The cells were transduced with empty vector, *Per1*, *Per2*, CBE (*Clock*, *Bmal1*, and empty vector), CBP1 (*Clock*, *Bmal1*, and *Per1*), and CBP2 (*Clock*, *Bmal1*, and *Per2*) before differentiation. *Igf2* KD cells and control cells were compared. (C) and (D) Differentiation index (C) and fusion index (D) of the cells used in (B). Data are presented as mean + SEM of biological triplicates. Each replicate includes n = 1,000-1,500 nuclei in (C) and (D). * p < 0.05, ** p < 0.01, and *** p < 0.001 with Student’s t-test compared with control KD cells in (A) and with empty vector cells in (B)-(D).

In a complementary study, *Per1* and *Per2* were overexpressed with retrovirus in C2C12 cells (Fig. S4 E) to investigate the effects on *Igf2* expression and differentiation. This study was done comparing control KD and *Igf2* KD cells to understand whether the effects of *Per* overexpression were dependent on the presence of *Igf2*. In the control KD cells, overexpressed *Per1* and *Per2* raised the *Igf2* mRNA level up to 4-fold higher than that with empty vector on differentiation days 3 and 5 (Fig. 3 B, *Igf2* in the Control KD columns). Because Per1 and Per2 regulate gene expression through binding to the Clock/Bmal1 complex, *Clock* and *Bmal1* were also overexpressed with or without *Per1* and *Per2*. However, the additional overexpression of *Clock* and *Bmal1* did not affect the *Igf2* level (Fig. 3 B, *Igf2* with CBE, CBP1, and CBP2 in the Control KD columns). A similar trend was observed in the effects of overexpressed *Per1* and *Per2* on differentiation. Whereas single overexpression of *Per1* or *Per2* raised the expression levels of differentiation-specific genes, differentiation index, and fusion index, an addition of *Clock* and *Bmal1* did not substantially alter the effects (Fig. 3, B-D). Importantly, all the effects by the overexpressed *Per1* and *Per2* were erased in *Igf2* KD cells (Fig. 3, B-D, *Igf2* KD1 columns), demonstrating that *Igf2* was an essential downstream effector for the overexpressed *Per1* and *Per2* to promote myoblast differentiation. Thus, up- and downregulation of *Per1* and *Per2* demonstrated the Per1/Per2-Igf2 axis as a novel myogenic pathway.

We also investigated whether Igf2 promotes muscle regeneration in an autocrine manner *in vivo* using satellite cell-specific conditional KO mice of *Igf2* (*Igf2* cKO). Satellite cell-specific cKO was particularly important for *Igf2* because the soluble factor from other tissues could regulate muscle regeneration. Although whole-body KO of *Igf2* impairs mouse growth, muscle-specific roles of *Igf2 in vivo* remained unclear. We disrupted only the paternally-inherited *Igf2* allele with *Pax7* promoter-driven CreERT2 because *Igf2* is imprinted with the paternal allele being preferentially expressed (Sasaki et al., 1992). After induction of CreERT2 with repeated tamoxifen injection, TA muscle was injured with barium chloride and muscle regeneration was compared with control mice that did not harbor the CreERT2 gene (Fig. 4 A). Depletion of the *Igf2* mRNA in satellite cells was verified by qPCR (Fig. 4 B). HE staining displayed that although *Igf2* cKO only slightly decreased myofiber size in uninjured muscle compared with control mice, the smaller size was more prominent in regenerating muscle on day 7 (Fig. 4, C-E). The size of eMHC(+) myofibers on day 4 was also smaller in *Igf2* cKO mice (Fig. 4, F and G). Although *Igf2* cKO did not increase the frequency of EdU(+)/MyoD(+) cells unlike *Per* KO, it decreased the population of EdU(-)/MyoD(+) cells, an indicator of delayed cell cycle exit in satellite cells by *Igf2* cKO (Fig. 4, H and I). Furthermore, differentiation index and fusion index in primary myoblasts were smaller in *Igf2* cKO mice compared with control mice *in vitro* (Fig. 4, J and K, and S4 F), verifying the previous results with C2C12 cells. Collectively, these findings indicated the requirement of autocrine Igf2 in myoblast differentiation and muscle regeneration.

**Figure 4.**
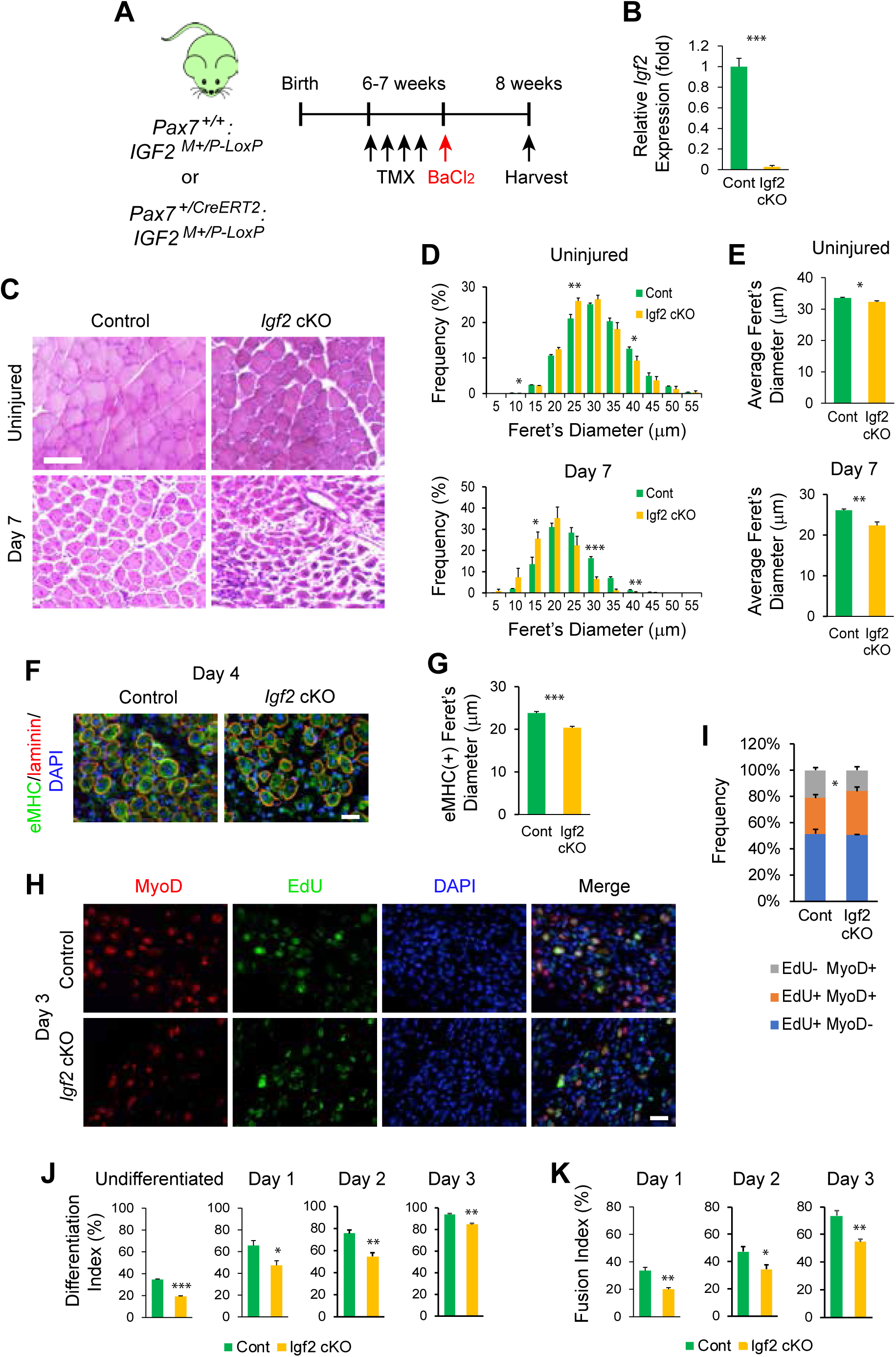
Impaired muscle regeneration in satellite cell-specific *Igf2* cKO mice. (A) Experimental scheme for satellite cell-specific *Igf2* cKO by injection of tamoxifen (TMX) and muscle regeneration from barium chloride (BaCl_2_)-induced injury. (B) Relative expression level of *Igf2* in isolated satellite cells comparing control and *Igf2* cKO cells. (C) HE staining of TA muscle sections prepared from uninjured and regeneration day 7 mice. TA muscle was injured with BaCl_2_ at ZT14 on day 0. Bar, 100 μm. (D) Size distribution of HE-stained myofibers containing centrally-located nuclei on the indicated time points. The minimal Feret’s diameter in each myofiber was measured. n = 3 for each genotype. (E) Average of the minimal Feret’s diameters of myofibers with centrally-located nuclei. (F) Immunofluorescence staining of TA muscle with antibodies against eMHC and laminin on day **1.** DNA was counterstained with DAPI. Bar, 25 μm. (G) Average of the minimal Feret’s diameters of eMHC(+) areas on day 4. n = 4 for each genotype. (H) Immunofluorescence staining of TA muscle sections with MyoD antibody and the EdU kit. Mice were injected with EdU on day 2 post-injury and TA muscle was harvested on day 3 for the staining. Bar, 25 μm. (I) Frequency of positive cells for EdU uptake and MyoD staining shown in (H). n = 3 for each genotype. (J) and (K) Differentiation index (J) and fusion index (K) of primary myoblasts prepared from control and *Igf2* cKO mice, representing biological triplicates with n = 80-110 nuclei in each replicate. Data are presented as mean + SEM. * p < 0.05, ** p < 0.01, and *** p < 0.001 with Student’s t-test compared with control.

### Circadian expression of *Igf2*

*Igf2* was likely to be expressed in a circadian manner as a target gene of Per1 and Per2. This possibility was tested by western blotting with C2C12 cells harvested every 4 hr after circadian synchronization with dexamethasone. The protein level of Bmal1 reached a peak at 44 hr after synchronization, which was anti-phasic to the expression patterns of Per1 and Per2 in control cells (Fig. 5 A). Igf2 expression reached peaks at 32-36 hr and 56 hr, similar to the patterns of Per1 and Per2. Phosphorylation of p38 (p-p38) followed the expression pattern of Igf2 as its downstream effector. In contrast, Igf2 and p-p38 were severely downregulated in *Per* KO cells. *Igf2* was also expressed in TA muscle in a circadian manner but the rhythms were largely lost in *Per1^-/-^* and *Per2^-/-^* mice (Fig. 5 B). These results demonstrate that *Igf2* shows circadian expression and that its expression level is regulated by Per1 and Per2.

**Figure 5.**
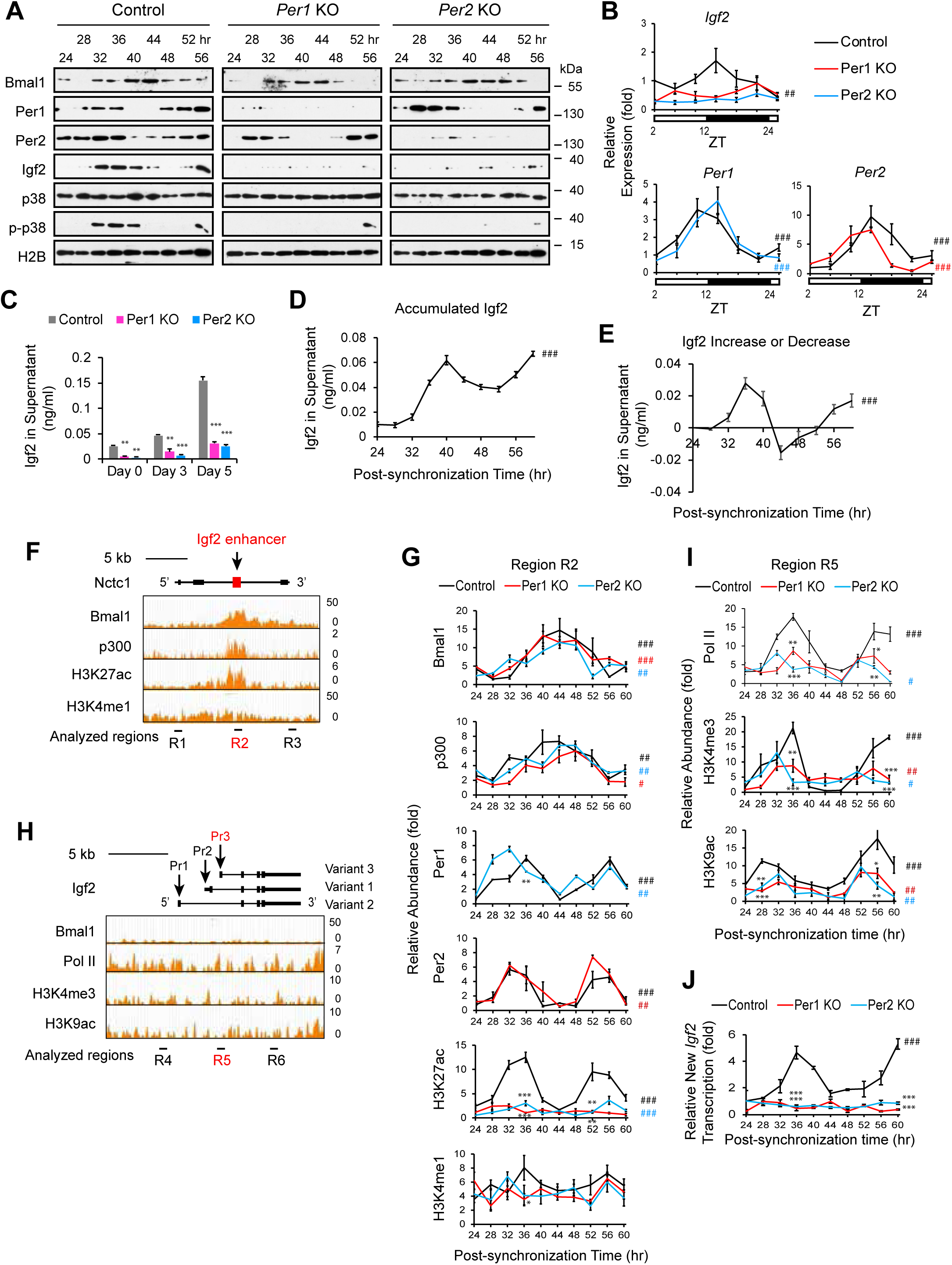
Circadian oscillation of the expression and epigenetics of the *Igf2* gene. (A) Western blotting with control and *Per* KO C2C12 cells harvested every 4 hr after circadian synchronization. Histone H2B was used as loading control. (B) Relative expression levels of *Igf2*, *Per1*, and *Per2* in TA muscle measured by qPCR. The value of WT mice at ZT2 was defined as 1.0. n = 3 mice with technical triplicates each. (C) Igf2 concentration in the C2C12 cell supernatant measured with ELISA. The culture medium was not replaced for 48 hr before measurement. (D) Igf2 concentration in the C2C12 cell supernatant after circadian synchronization. Cells were treated with dexamethasone between -1 and 0 hr for synchronization. The culture medium was replaced with fresh growth medium at 0 hr and was not changed until the harvest at the indicated time point. The concentration indicates the accumulated Igf2 in the medium. (E) The change of the Igf2 concentration in (D) was highlighted by displaying the change of the concentration between two time points. (F) ChIP-seq analyses of the *Igf2* enhancer within the *Nctc1* gene downloaded from the Gene Expression Omnibus (GEO) database. See methods for the accession number of each data set. R1-R3 indicate the regions amplified by PCR in (G) and Figure S5 A. (G) ChIP-PCR analyses of indicated proteins in control and *Per* KO C2C12 cells at the region R2. Relative abundance compared with control IgG is shown. Peak values of control cells that are higher than those of *Per1* KO and *Per2* KO cells are highlighted with asterisks. (H) ChIP-seq analyses of the *Igf2* promoters downloaded from the GEO database. See methods for the accession number of each data set. R4-R6 indicate the regions amplified by PCR in (I) and Fig. S5 D. Variant 1, 2, and 3, correspond to NM_010514, NM_001122736, and NM_001122737, respectively. Pr1 to Pr3 indicate Promoter 1 to Promoter 3. (I) ChIP-PCR analyses of indicated proteins in control and *Per* KO C2C12 cells at the R5 region. (J) Nascent transcript analysis with a nuclear run-on assay comparing control and *Per* KO cells. Synchronized C2C12 cells were labeled with 5-ethynyl uridine (EU) for 4 hr before harvesting every 4 hr and EU(+) RNA was isolated with a kit, followed by RT-PCR of the indicated genes. The values were normalized against *Clock* mRNA, whose expression was not influenced by circadian rhythms. The value with control cells at 24 hr post-synchronization was defined as 1.0. Data are presented as mean ± SEM of biological triplicates. * p < 0.05, ** p < 0.01, and *** p < 0.001 with Student’s t-test compared with control. ^#^ 24 h rhythmicity with Cosinor (p < 0.05), ^##^ P < 0.01, and ^###^ P < 0.001.

Next, the concentration of Igf2 in the culture supernatant of C2C12 cells was measured with ELISA. The concentration was approximately 0.025 ng/ml with undifferentiated cells and was increased 6-fold during differentiation as previously reported (Fig. 5 C) (Florini et al., 1991). Although Igf2 in the supernatant of *Per* KO cells was also increased, the level remained less than 20% of the control level on day 5, consistent with PCR and western blotting results. The Igf2 concentration with control cells also displayed oscillation that was similar to the western blotting result (Fig. 5 D). The concentration represented the amount of accumulated Igf2 since 0 hr, when dexamethasone was replaced with fresh culture medium. The result likely reflected the gain by secretion and the loss by degradation and attachment to the culture dish and cell surface. The oscillation became more evident when an increase or a decrease between two time points were plotted (Fig. 5 E).

We also examined whether exogenous Igf2 could rescue the disrupted differentiation of *Per* KO cells by adding Igf2 to the culture medium from day 0 onward. Differentiation index and fusion index on days 3 and 5 exhibited improved differentiation with 1 ng/ml Igf2, reaching 40- 60% levels of those observed with control cells without exogenous Igf2 (Fig. S4 G). Activation of *Ckm* and *Myog* by Igf2 was more efficient; the mRNA levels became indistinguishable from those with control cells without exogenous Igf2 on day 3 for both genes and on day 5 for *Myg* (Fig. S4 H). Thus, exogenous Igf2 could substantially rescue the disrupted differentiation of *Per* KO cells, consistent with the notion of the Per1/Per2-Igf2 axis. Note that the concentration of effective Igf2 in the culture medium in these experiments was unknown due to the presence of IGFBPs.

### Epigenetic regulation of the *Igf2* expression by Per1 and Per2

To elucidate how Per1 and Per2 promoted *Igf2* expression, epigenetic changes caused by *Per* KO were studied with ChIP-qPCR. An *Igf2* enhancer containing two E-boxes is embedded within an intron of the *Nctc1* gene located 105 kb downstream of the *Igf2* promoter (Alzhanov and Rotwein, 2016; Alzhanov et al., 2010). Publicly available ChIP-seq data obtained with non-synchronized myoblasts demonstrated binding peaks of Bmal1, its binding partner histone acetylase p300 (Papazyan et al., 2016; Takahashi, 2017b), and an enhancer marker H3K27ac (acetylated lysine 27 in histone H3) at the *Igf2* enhancer, whereas another enhancer marker H3K4me1 was broadly distributed (Fig. 5 F). Our ChIP-qPCR with synchronized control C2C12 cells demonstrated circadian binding of Bmal1 and p300 at the enhancer and the temporal profile was antiphasic to those of Per1 and Per2 as anticipated (Fig. 5 G). The binding pattern of Bmal1 was not significantly altered by KO of *Per1* or *Per2*. The abundance of H3K27ac also demonstrated an oscillatory pattern that was flattened in *Per1* and *Per2* KO cells. The level of H3K4me1 did not exhibit a clear circadian pattern in control or *Per* KO cells. The histone modification patterns were consistent with the general trend that whereas H3K27ac marks active enhancers, H3K4me1 is present in both active and inactive/poised enhancers (Creyghton et al., 2010; Rada-Iglesias et al., 2011). All these protein levels were substantially decreased outside the enhancer except for the less prominent or no decrease of H3K4me1 (Fig. S5 A, Regions R1 and R3), underscoring the enhancer specificity of our results.

Muscle cells primarily use Promoter 3 among the three promoters of the *Igf2* gene (Baral and Rotwein, 2019; Kou and Rotwein, 1993), which was verified by qPCR (Fig. S5, B and C). Downloaded ChIP-seq data with non-synchronized myoblasts did not show a specific increase or decrease of Bmal1, RNA polymerase II (Pol II), or histone markers for active genes (H3K4me3 and H3K9ac) at Promoter 3 (Fig. 5 H, Region R5). However, synchronized C2C12 cells again demonstrated a Per-dependent increase of these proteins at Promoter 3 (Fig. 5 I) with weaker or no oscillation in the surrounding regions (Fig. S5 D for Regions R4 and R6). Thus, both Per1 and Per2 are necessary for the circadian dynamics of multiple epigenetic markers characteristic of gene activation at the *Igf2* enhancer and Promoter 3.

To understand the functional significance of the circadian epigenetics, the temporal profile of nascent *Igf2* mRNA was quantified with synchronized cells. A nuclear run-on assay demonstrated that nascent *Igf2* mRNA was most abundant at time points when the *Igf2* enhancer and promoter were enriched with active gene markers in control cells (Fig. 5 J, 36 hr and 60 hr). However, the *Igf2* level remained low throughout the process with *Per* KO cells as expected. Therefore, the circadian transcriptional changes of *Igf2* indeed reflected the epigenetic dynamics of the gene.

The interaction between the *Igf2* enhancer and Promoter 3 has been shown in differentiating myoblasts and skeletal muscle (Alzhanov et al., 2010; Yoon et al., 2007). Because chromatin loop formation is also regulated by circadian rhythms (Pacheco-Bernal et al., 2019; Yeung et al., 2018), we hypothesized that the *Igf2* enhancer-promoter interaction would also demonstrate circadian oscillation. This possibility was examined with Chromosome Conformation Capture (3C) by studying the interaction between Promoter 3 (anchor point in 3C) and the enhancer at 24 and 36 hr post-synchronization, which corresponded to the nadir and the peak of the *Igf2* level, respectively (Fig. 6 A). The interaction (crosslinking frequency) at 36 hr was more than twice as high as it was at 24 hr in control cells (Fig. 6 B). This pattern was preserved in *Per* KO cells but the crosslinking efficiency became less than half compared with control cells (Fig. 6, B and C). Therefore, both Per1 and Per2 are required for the circadian dynamics of the promoter-enhancer interaction at the *Igf2* gene, just like the epigenetic modifications.

**Figure 6.**
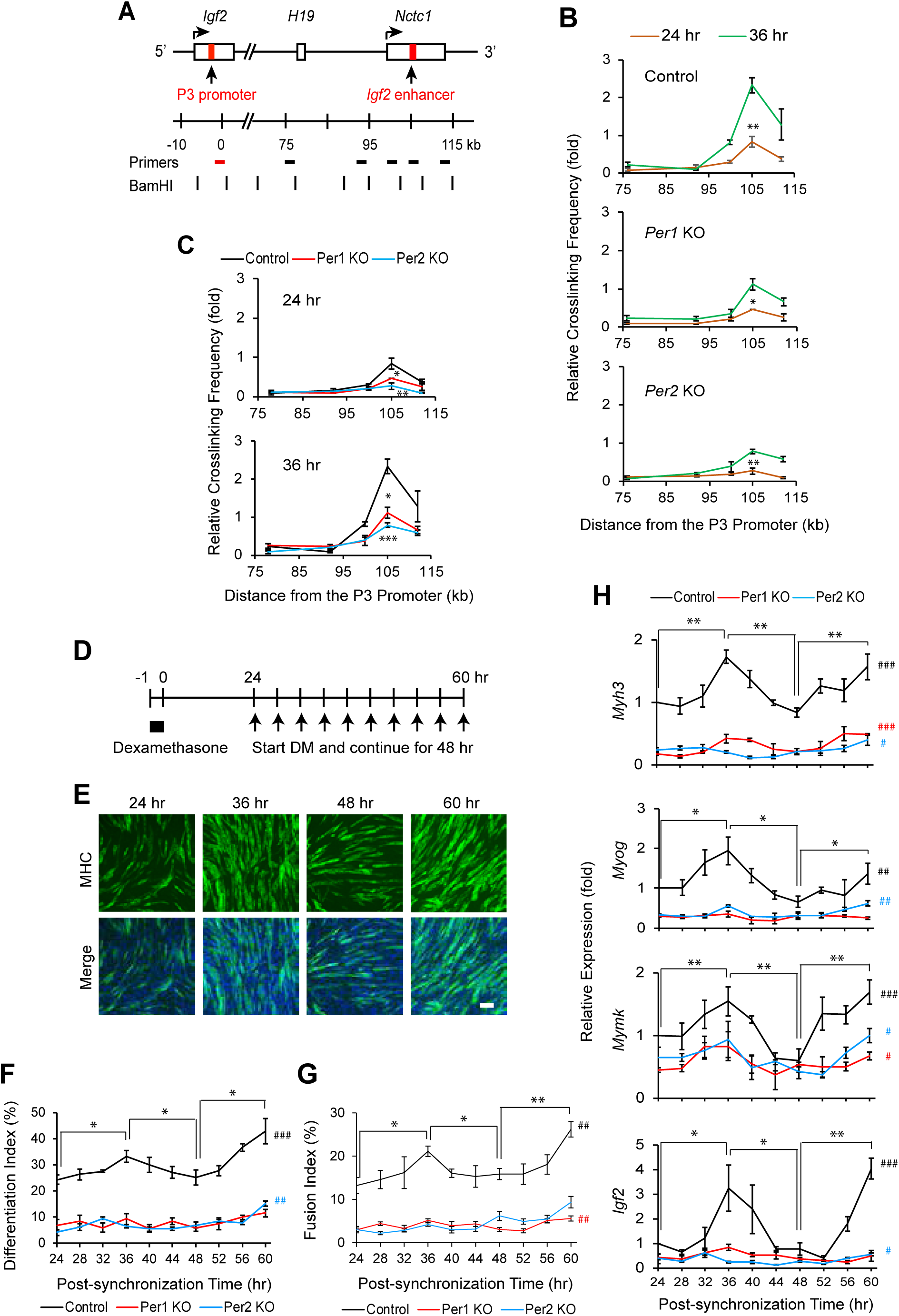
Circadian regulation of the *Igf2* gene and C2C12 cell differentiation. (A) Locations of the primers used in the 3C experiments and BamHI sites in relation to the *Igf2* Promoter 3 and enhancer. The primer shown in red was used in combination with one of the primers shown in black in 3C and the results were plotted in (B) and (C). (B) and (C) Relative crosslinking frequency obtained with 3C comparing different time points (B) and *Per* KO cells and control (C). The value obtained with the *Clock* gene primers was defined as 1.0. (D) Schedule of circadian synchronization and initiation of differentiation. After incubation with dexamethasone between -1 hr and 0 hr, the culture medium was replaced with growth medium containing 10% fetal bovine serum at 0 hr. The culture medium was replaced with differentiation medium (DM) containing 5% horse serum at different time points every 4 hr (arrows). Differentiation was continued for 48 hr from each starting point before fixation or harvest for various analyses. (E) Immunofluorescence staining C2C12 cells with MHC antibody and Hoechst 48 hr after starting differentiation at indicated time points shown in (D). Bar, 200 μm. (F)-(H) Analyses of differentiation index (F), fusion index (G), and relative expression of differentiation-specific genes (H) with C2C12 cells that were induced to differentiate at the indicated post-synchronization time points. * p < 0.05, ** p < 0.01, and *** p < 0.001 with Student’s t-test comparing two time points in (B), control and *Per* KO cells in (C), and different time points in control cells in (F)-(H). Graphs show mean ± SEM of biological triplicates. Each replicate includes n = 1,000-1,500 nuclei in (F) and (G). ^#^ 24 h rhythmicity with Cosinor (p < 0.05), ^##^ P < 0.01, and ^###^ P < 0.001.

### Coupling of differentiation efficiency and the circadian timing of differentiation initiation

The above findings led us to a hypothesis that myoblasts could differentiate more efficiently if differentiation cues are provided at the circadian timing when *Per1*, *Per2*, and *Igf2* are highly expressed compared with other time points (“primed”). This was evaluated with C2C12 cells that were induced to differentiate at different time points after synchronization. Indeed, the cells differentiated more efficiently when induced at 36 hr and 60 hr post-synchronization compared with 24 hr and 48 hr as demonstrated by higher differentiation index, fusion index, and the expression levels of differentiation-specific genes (*Myh3*, *Myog*, and *Ckm*) and *Igf2* after 48 hr of differentiation (Fig. 6, D-H). This trend was in agreement with the epigenetic modifications of the *Igf2* enhancer and promoter. Note that since the cells kept proliferation between 24 hr and 60 hr before the differentiation induction, the increased cell density *per se* could promote differentiation in the later phase. In this sense, the drop of the differentiation efficiency at 44-48 hr in comparison to 36 hr was more significant than the increased differentiation from 48 hr to 60 hr. *Igf2* KD cells differentiated poorly regardless of when differentiation was initiated (Fig. S5, E-G).

Since *Per1*, *Per2*, and *Igf2* were also expressed in TA muscle in a circadian manner (Fig. 5 B), the timing of the injury could be an important factor for muscle regeneration. To assess this possibility, TA muscle was injured at ZT2 (low *Per1*, *Per2*, and *Igf2*; the early inactive phase of mice) and ZT14 (high *Per1*, *Per2*, and *Igf2*; the early active phase) to compare the regeneration efficiency. HE staining clearly showed larger TA muscle in the ZT14 WT mice than in the ZT2 mice on day 4.5 but this difference was lost by day 14 (Fig. 7, A and B, and S5 H). There was no statistically significant difference between ZT2 and ZT14 injuries in the single and double KO mice of *Per*1 and *Per2*. The average diameter of eMHC(+) myofibers was also longer in WT mice damaged at ZT14 than that in the ZT2 damage and this difference was also lost in the single and double KO mice (Fig. 7, C and D). Moreover, the frequency of the EdU(+)/MyoD(+) population was diminished in the ZT14 WT mice compared with ZT2 mice on day 4.5 and 5.5, suggesting early cell cycle exit (Fig. 7, E and F). Finally, fibrosis was also less abundant in the ZT14 WT mice than in the ZT2 mice on day 14; this difference was again erased in the single and double KO mice (Fig. 7, G and H). These results collectively indicate that circadian timing of injury affects the efficiency of TA muscle regeneration in a *Per1*- and *Per2*-dependent manner.

**Figure 7.**
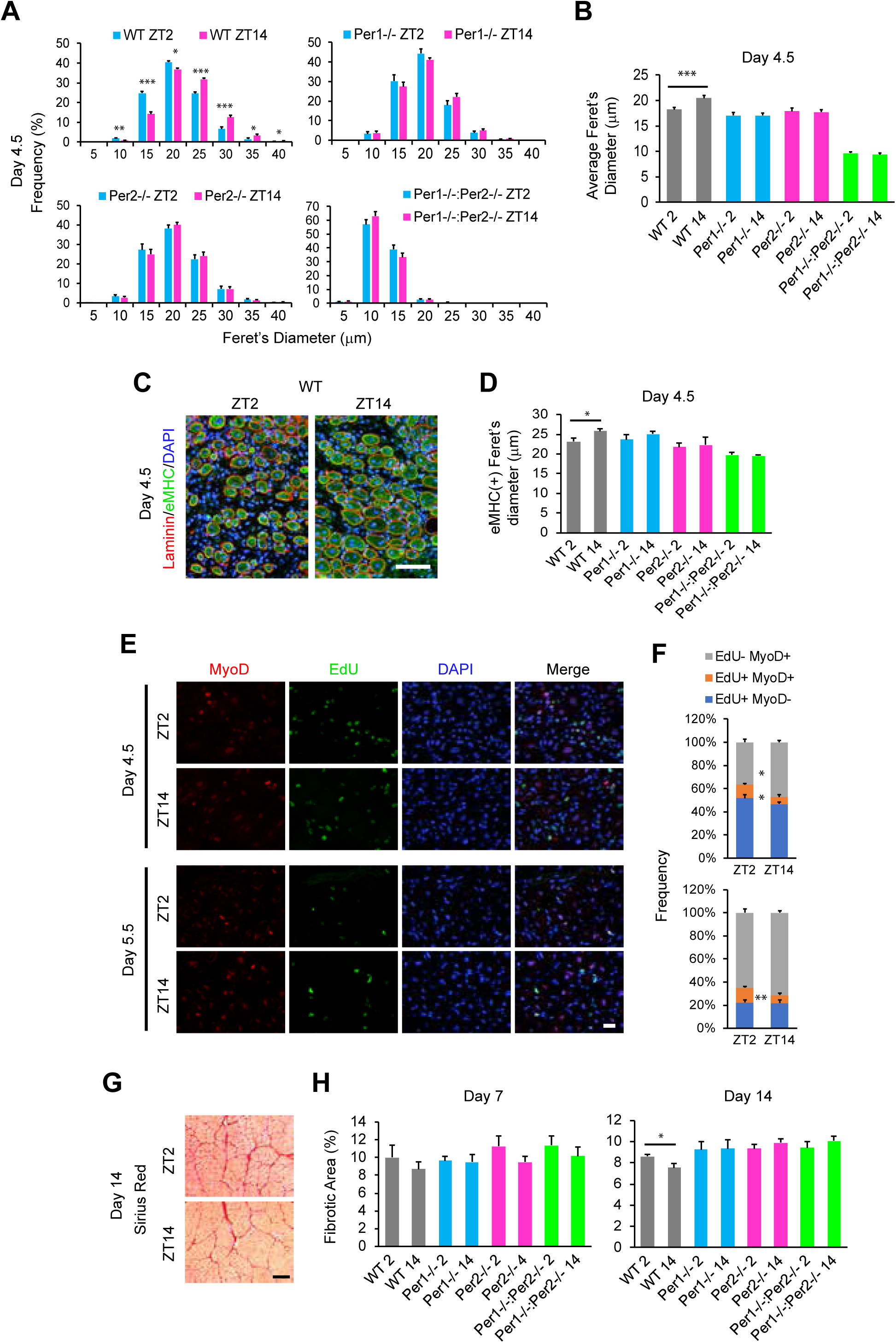
Differential regeneration efficiency of TA muscle depending on the circadian timing of the injury. (A) Size distribution of HE-stained myofibers containing centrally-located nuclei on day 4.5. TA muscle was injured with barium chloride at ZT2 or ZT14 on day 0 and harvested on day 4.5. The minimal Feret’s diameter of each myofiber was calculated. n = 8 mice with 4 males and 4 females in each group in (A) and (B). (B) Average of the minimal Feret’s diameters of myofibers with centrally-located nuclei on day 4.5. 2 and 14 at the end of each genotype indicate the injury time at ZT2 and ZT14, respectively. (C) Immunofluorescence staining of WT TA muscle injured at ZT2 and ZT14 with antibodies against eMHC and laminin on day 4.5. DNA was counterstained with DAPI. Bar, 100 μm. (D) Average of the minimal Feret’s diameters of the eMHC(+) areas on day 4.5. n = 4 mice. (E) Immunofluorescence staining of WT TA muscle sections with the MyoD antibody and an EdU kit. TA muscle was injured with barium chloride at ZT2 or ZT14 on day 0 and EdU was intraperitoneally injected 96 or 120 hr later. The muscle was harvested 12 hr later as day 4.5 or 5.5. Bar, 25 μm. (F) Frequency of positive cells for EdU uptake and MyoD staining in TA muscle sections shown in (E). n = 4 mice. (G) Sirius red staining of WT TA muscle on day 14 post-injury. Bar, 200 μm. (H) The area percentage of fibrosis indicated by positive Sirius red staining on days 7 and 14. Data are presented as mean + SEM. * p < 0.05, ** p < 0.01, and *** p < 0.001 with Student’s t-test. The values at ZT14 in Fig. 1 were reused in these figures.

## Discussion

The most novel message of the present work is that the efficiency of myoblast differentiation and muscle regeneration is dependent on the circadian timing when these events are triggered. As a mechanistic explanation obtained with the myoblast model, the *Igf2* gene was primed toward activation in a circadian manner while the cells were still in the proliferation medium. Despite extensive studies of circadian regulation of cell proliferation and differentiation (Benitah and Welz, 2020; Dierickx et al., 2018; Paatela et al., 2019), studies focused on the mechanistic influence of circadian timing on tissue regeneration are quite limited. One of the few studies concerns fibroblast migration during skin wound healing (Hoyle et al., 2017). Fibroblast mobilization to a mouse skin incision site, an early and essential step in wound healing, was greater when the wound was inflicted at ZT13 than at ZT5. Additionally, when a skin explant was harvested and wounded at different time points, the number and volume of fibroblasts invading the wound area were higher in the explant harvested at ZT13 than at ZT5. Circadian regulation of actin polymerization, which controls migration and adhesion, is one of the mechanisms for the time-dependent difference. In a related phenomenon, circadian timing of physical exercise influences muscle strength and oxidative capacity (Wolff and Esser, 2019). For example, muscle atrophy in the mouse hind limb due to reduced gravity was prevented more effectively by intermittent weight bearing between ZT12-ZT16 than between ZT20-ZT0 (Aoyama et al., 2018). Although we linked circadian timing and the efficiency of muscle regeneration from an acute injury, it remains an open question whether chronic muscle damages, such as exercise-induced injuries, exhibit the same level of circadian responses. At the mechanistic level, we demonstrated that Igf2 is a required downstream effector of Per1/Per2 for myogenesis and that Igf2 is expressed in a circadian manner. However, we did not determine whether it was the overall expression level or rhythmicity of Pe1/Per2 and Igf2 that is important for myogenesis and muscle regeneration. Experiments with KO cells and mice cannot distinguish the two possibilities. A recently reported artificial circadian clock in clock-deficient mice could provide a valuable model to directly address this question (D’Alessandro et al., 2015).

Our findings on the circadian timing-dependent differentiation and regeneration should be interpreted in a broader perspective of circadian metabolic regulation that defines the availability of energy and cellular building blocks (Aoyama and Shibata, 2017; Aoyama and Shibata, 2020; Panda, 2016). A circadian transcriptome analysis of muscle uncovered clustered expression of genes with a common metabolic function at specific circadian phases in the mouse under constant darkness with *ad libitum* feeding (Hodge et al., 2015). Specifically, the genes involved in carbohydrate catabolism (the early active/dark phase), carbohydrate storage (the mid-active/dark), lipogenesis (the end of the active/dark phase), and fatty-acid uptake and β-oxidation (the mid-inactive/light phase) reached peaks at distinct circadian phases as indicated in the parentheses. Metabolomic profiling of muscle also demonstrated neutral lipid storage and decreased lipid and protein catabolism in the late inactive phase (Dyar et al., 2018). Given the global circadian oscillation of the numerous metabolites essential for tissue turnover, circadian timing could create a preferred time window for an effective response to major tissue disruption and repair although experimental evidence is lacking. The interaction between the Per1/Per2-Igf2 axis and the global metabolic oscillation awaits further studies. This is further supported by a recent study demonstrating regulation of the Per protein level by insulin and Igf1, which are regulated by feeding and share signaling pathways with Igf2, at the transcription and translation levels (Crosby et al., 2019).

Per1 and Per2 are generally considered as transcription repressors. In our study, however, more than 1,000 genes were commonly upregulated by *Per 1* KO and *Per2* KO cells, suggesting that Per1 and Per2 can act as direct or indirect gene activators in this context. Per-induced gene activation has been demonstrated with several genes involved in sodium channels in the kidney (Gumz et al., 2009; Richards et al., 2014; Stow et al., 2012). In addition, Bmal1 and Per1 are required for the circadian activation of prolactin in a rat mammotrope cell line (Bose and Boockfor, 2010). Moreover, Per2 activates *Cry1* by removing the Clock/Bmal1/Cry1 repressor complex from the *Cry1* promoter in an ectopic expression model (Chiou et al., 2016). This study also showed that genes with complex promoters can be repressed or de-repressed by Per, depending on the regulatory elements at the promoters. Furthermore, Per2 can activate its own promoter, which has been explained by the blocking of the binding of Cry to the Clock/Bmal1 complex (Akashi et al., 2014). In our study, although the Per proteins bind the *Igf2* enhancer, it remains unknown whether they directly contribute to the transcription activation of *Igf2*. Because Per proteins bind thousands of loci along with Clock/Bmal1, *Igf2* activation could be an indirect activity of Pers. Identification of the binding proteins of Per1 and Per2 would be an important next step to further clarify how Per1 and Per2 activate *Igf2* during myoblast differentiation.

This study revealed circadian regulation of myoblast differentiation and muscle regeneration and demonstrated epigenetic regulation of the *Igf2* gene by Per1 and Per2 as one of the underlying mechanisms using a myoblast differentiation model. Future genome-wide epigenetic analysis of histone modifications and chromatin interactions would further uncover other unexpected underpinnings for the time-of-the-day-dependent regeneration of muscle and other tissues.

## Materials and Methods

### *Per* Knockout Mice

All protocols were approved by the Institutional Animal Care and Usage Committee of the University of Minnesota (1902-36737A and 1903-36906A). *Per1^+/-^* mice (B6.129-*Per1^tm1Drw^*/J, stock # 010491) and *Per2^+/-^* mice (B6.129-*Per2^tm1Drw^*/J, stock# 010492) were purchased from Jackson Laboratory. *Per1^-/-^*, *Per2^-/-^*, *Per1^-/-^*:*Per2^-/-^*, and wild type (WT) mice were obtained by breeding and identified by genotyping according to Jackson Laboratory protocols. The same number of male and female mice at the age of 8-10 weeks old were mixed in each group. The number of used mice are described in each figure legend. Mice were entrained at 12 hr-light and 12 hr-dark cycles (6:00-18:00 light and 18:00-6:00 dark) for two weeks before experiments. ZT0 corresponds to 6:00 and ZT12, 18:00. TA muscle was injected with 50 μl 1.2% BaCl_2_ in 0.9% NaCl at ZT2 or ZT14 to induce muscle injury and subsequent regeneration. Mice were euthanized on day 4.5, 7, and 14 after injury and the TA muscle was extracted at ZT2 or ZT14. In addition, uninjured TA muscle was isolated every 4 hr for qPCR. Mice were monitored by the Research Animal Resources staff of the University of Minnesota in specific pathogen-free housing. Mice were given standard chow and access to drinking water without restrictions. Mice were euthanized via CO_2_ inhalation. All methods align with the Panel of Euthanasia of the American Veterinary Medical Association recommendations.

### Conditional *Igf2* knockout mice

*Pax7^CreERT2^* mice (B6.Cg-*Pax7^tm1(cre/ERT2)Gaka^*/J, stock # 017763) and *IGF2^LoxP/LoxP^* mice (*Igf2^tm1.1Thor^*/J, stock # 032493) were purchased from Jackson Laboratory (Modi et al., 2015; Murphy et al., 2011). *Igf2* is imprinted with preferential expression of the paternal allele (Sasaki et al., 1992). Superscript (P) and (M) indicate paternal and maternal inheritance of the *Igf2* allele in the *IGF2 ^M-LoxP/P-LoxP^* mice, respectively. *IGF2^M-LoxP/P-LoxP^* male mice were bred with *Pax7^+/CreERT2^* female mice to generate *Pax7^+/+^:IGF2^M+/P-LoxP^* (control) and *Pax7^+/CreERT2^:IGF2^M+/P- LoxP^* (conditional knockout) mice. Genotyping to detect the mutant alleles was performed by PCR using the primers shown in Table S1. Cre recombination was induced with tamoxifen (TMX) (T5648, MilliporeSigma) at 75 μg/g body weight 4 times between 6 and 7 weeks of age. TA muscle was injured with BaCl_2_ at 7 weeks of age and the muscle was extracted within one week. Control mice that contained the WT *Pax7* allele were also injected with TMX.

### Preparation of Primary Myoblasts

Mouse muscle mononuclear cells were prepared from hind limbs of 8-10 weeks old male mice as previously described (Asakura et al., 2002; Motohashi et al., 2014). More specifically, muscles were minced and digested with 0.2% collagenase type 2 (Worthington, CLS-2) in Dulbecco’s Modified Eagle Medium (DMEM) to dissociate muscle cells. Satellite cells were purified with LD columns (Miltenyi Biotec, 130-042-901) by negative selection with antibodies against CD31-PE, CD45-PE, and Sca1-PE, followed by anti-PE MicroBeads (Miltenyi Biotec, 130-048-801). See Table S2 for the details of the antibodies. Positive selection was subsequently applied with an antibody against biotin-conjugated integrin α7-biotin and anti-biotin MicroBeads (Miltenyi Biotec, 130-090-485), followed by MS columns (Miltenyi Biotec, 130-042-201). Isolated satellite cells were cultured on dishes coated with 0.01% rat tail collagen (BD Biosciences, 354236) in myoblast growth medium (HAM’s F-10 medium with 20% FBS, 10 ng/ml basic fibroblast growth factor (Thermo Fisher Scientific, PHG0263), penicillin (100 U/ml), and streptomycin (100 mg/ml)) at 37°C with 5% CO_2_. Low-passage satellite cell-derived primary myoblasts (typically less than eight passages) were used for differentiation and immunostaining. DMEM with 5% horse serum (HS), penicillin, and streptomycin was used for myogenic differentiation.

### Culture of C2C12 cells

Female mouse myoblast C2C12 cells (American Type Culture Collection, CRL-1772) were maintained in the growth medium (10% fetal bovine serum (FBS) in DMEM) in a 37°C and 5% CO_2_ incubator. Cells were authenticated by immunofluorescence staining of MyoD. Differentiation was induced on day 0 when cells were at 90% confluence by rinsing with phosphate-buffered saline (PBS) twice and adding the differentiation medium (5% HS in DMEM). The medium was changed every two days thereafter. Concentration of endogenous Igf2 was measured with a mouse Igf2 ELISA kit (Thermo Fisher Scientific, EMIGF2). Because of the low Igf2 concentration, 300 μl culture supernatant was added seven times for 3 hr each (2.1 ml in total). The effect of exogenous Igf2 was tested with recombinant human IGF2 (Peprotech, 100-12). To synchronize circadian rhythms, C2C12 cells were seeded on day -1 and 0.2 μM dexamethasone was added at -1 hr on day 0. Cells were washed with PBS twice and fresh 10% FBS in DMEM was added at 0 hr. Cells were harvested for PCR or fixed for immunofluorescence staining every 4 hr at indicated time points.

### Hematoxylin Eosin (HE) Staining

Cryosections of the TA muscle were fixed with 2% paraformaldehyde for 10 min. The sections were then treated as follows: deionized water for 1 min, Harris Modified Hematoxylin (Thermo Fisher Scientific, SH26-500D) for 2 min, tap water for 1 min, deionized water for 1 min, Eosin-Y (Thermo Fisher Scientific, 22-220-104) for 5 min, 95% ethanol for 30 sec, 100% ethanol for 2 min twice, and xylene for 10 min twice. The sections were mounted with Permount (Thermo Fisher Scientific, SP15-100). Images were captured with the cellSens Entry 1.11 software (Olympus) and a DP26 camera (Olympus) attached to an IX73 microscope with a UPlan FL N 10X/0.30 Ph1 lens (Olympus). The minimal Feret’s diameter of each myofiber was quantified with Fiji (NIH).

### Sirius Red Staining

The Sirius red solution was composed of 1% Direct Red 80 (MilliporeSigma, 365548) in 1.3% picric acid. TA sections were fixed with acetone, pre-chilled at -20°C, for 10 min. The sections were then washed in deionized water for 1 min, stained with Sirius red for 15 min, and rinsed with 0.5% acetic acid for 1 min. The sections were subsequently washed with 100% ethanol for 2 min and twice with xylene for 10 min. Finally, the sections were mounted with Permount for taking images as described for HE staining. Quantification of Sirius red (+) fibrosis was done using the entire section area with Fiji (Shimizu-Motohashi et al., 2015).

### Immunofluorescence Staining of TA Sections

Sections were fixed with 2% paraformaldehyde for 10 min, followed by permeabilization with 0.2% Triton X-100 in PBS for 5 min. The sections were treated with two blocking reagents: 3% Mouse-on-Mouse Blocking Reagent (Vector, MKB-2213) in PBS for 1 hr and 10% bovine serum albumin (BSA) in PBS for 30 min. Primary antibodies against eMHC and laminin diluted in 10% BSA in PBS were applied overnight, followed by washing twice with 0.01% Triton-X100 in PBS. The secondary antibodies Alexa Fluor 488 donkey anti-mouse IgG and Alexa Fluor 594 donkey anti-rat IgG diluted in 10% BSA in PBS were used for 1 hr. DNA was counterstained with 4’,6’-diamidine-2’-phenylindole dihydrochloride (DAPI, MilliporeSigma, 10236276001). Sections were mounted using Fluorescent Mounting Medium (DAKO, S302380-2). To label proliferating myoblasts, mice were intraperitoneally injected with 5-ethynyl-2’-deoxyuridine (EdU) at 50 μg/g body weight on day 4 and euthanized 12 hr later on day 4.5 post-injury. EdU was detected using Click-iT EdU Alexa Fluor 488 Imaging Kit (ThermoFisher Scientific, C10337) followed by staining with anti-MyoD antibody, Alexa Fluor 594 donkey anti-rabbit IgG antibody, and DAPI (Verma et al., 2018). Fluorescence images were captured using the Metamorph Basic software (Molecular Devices) and an ORCA-flash4.0LT camera (Hamamatsu) attached to an IX73 microscope (Olympus) with an LUCPlan FL N 20x/0.45 Ph1 lens. Images were processed with Photoshop and Illustrator CS6 (Adobe). The minimal Feret’s diameter of each myofiber was quantified with Fiji (NIH).

### Gene Knockdown in C2C12 cells

On day 1, 293FT cells (Thermo Fisher Scientific, R70007) were seeded in DMEM with 10% FBS at 3×10^5^ cells/well in a 12-well plate. On day 2, cells were transfected with 0.5 μg pLKO.1 lentivirus vector encoding an shRNA sequence shown in Table S3, along with 0.2 μg each of pCMV-VSV-G (Addgene, 8454), pRSV-Rev (Addgene, 12253), and pMDLg/pRRE (Addgene, 12251) using 2.75 μl Lipofectamine 2000 (Thermo Fisher Scientific, 11668019). The culture medium was replaced with fresh DMEM with 10% FBS 5 hr after transfection. On day 4, C2C12 cells were seeded at 1×10^5^ cells/well in 12-well plates. On day 5, culture supernatant of the transfected 293FT cells was applied to a 0.45 μm syringe filter and added to C2C12 cells with 0.8 μg/ml polybrene (MilliporeSigma, H9268). The culture medium was replaced with fresh DMEM with 10% FBS on day 6. Virus-integrated cells were selected with 1 μg/ml puromycin dihydrochloride (MB Bio, 100552) between days 7 and 14. Selected cells were expanded and frozen in liquid nitrogen.

### Gene Knockout in C2C12 cells

On day 1, C2C12 cells were seeded in DMEM with 10% FBS at 1.8×10^5^ cells/well in 12-well plates. Cells were transfected with 0.5 μg single guide RNA (sgRNA, Synthego) against *Per1* and *Per2* (Table S3), 0.5 μg CleanCap Cas9 mRNA (TriLink, L-7206), 0.2 μg CleanCap mCherry mRNA (TrLink, L-7203), and 1 μl TransIT (Mirus Bio TransIT-mRNA transfection kit, MIR-2225). More than 90% of the cells were mCherry (+) with fluorescence microscope observation on day 2. Cells were subcultured on day 3 to expand and freeze. A small aliquot of cells was used for Sanger sequencing of genomic DNA to verify the knockout efficiency.

### Overexpression of *Clock*, *Bmal1*, *Per1*, and *Per2*

Mouse *Clock*, *Bmal1*, *Per1*, and *Per2* cDNAs were inserted into the pMXs-IP vector (Kitamura et al., 2003). PLAT-E cells (Morita et al., 2000) were seeded at a density of 2.5×10^5^ cells/well in a 12-well plate with 10% FBS on day 1. On day 2, cells were transfected with 750 ng of the pMXs-IP vectors with 2.3 μl FuGENE 6 (Promega, E2691). The culture medium was replaced with fresh DMEM with 10% FBS and C2C12 cells were seeded at a concentration of 1×10^5^ cells/well in a 12-well plate on day 3. PLAT-E cell medium containing the virus was harvested on day 4 and filtered through a 0.45 μm syringe filter before transduction into C2C12 cells with 8 μg/ml polybrene (MilliporeSigma, H9268). The medium was replaced again on day 4 prior to the second transduction on day 5. Starting on day 6, 1 μg/ml puromycin dihydrochloride selection for virus-integrated cells was performed for 5 days. Proliferating cells were frozen in liquid nitrogen or used in experimentation.

### Immunofluorescence Staining of Cells

Primary myoblasts were fixed with 2% paraformaldehyde for 10 min and blocked with 1% BSA in PBS for 30 min. Permeabilized cells were stained with antibodies against MHC and MyoD and then incubated with secondary antibody Alexa Fluor 488 anti-mouse IgG and Alexa Fluor 594 anti-rabbit IgG. DNA was counterstained with DAPI. To measure proliferating primary myoblasts, cells were pulsed with 1 μg/ml of EdU 3 hr before harvest. EdU was detected using a Click-iT EdU Alexa Fluor 488 Imaging Kit (ThermoFisher Scientific, C10337) followed by staining with anti-MyoD antibody, Alexa Fluor 594 donkey anti-rabbit IgG antibody, and DAPI (Verma et al., 2018). Fluorescence images were captured using Metamorph Basic software (Molecular Devices) with LUCPlanFLN 20x or 10x objective lens (Olympus) with 0.45 Ph1 aperture and a C11440-42U digital camera (Hamamatsu) attached to an IX73P2F microscope (Olympus). The images were processed with Adobe Photoshop and Illustrator CS6. Differentiation index was defined as a percentage of nuclei (DAPI-stained structure) existing within MHC(+) cells. Fusion index is a percentage of nuclei that were located in MHC(+) cells containing more than one nuclei. Differentiation index, fusion index, and EdU uptake were analyzed with biological triplicates, including 80-130 nuclei in each replicate.

C2C12 cells were similarly stained with primary antibody against MHC, secondary antibody Alexa Fluor 488 goat anti-mouse IgG, and 5 μg/ml Hoechst 33342 (MilliporeSigma, B2261). Differentiation index, fusion index, and EdU uptake were analyzed with biological triplicates, including 1,000-1,500 nuclei in each replicate.

### Western Blotting

Whole-cell extracts obtained from 2×10^5^ cells with an NE-PER Nuclear and Cytoplasmic Extraction kit (Thermo Fisher Scientific, 78833) were loaded into a 12% SDS-PAGE gel. After completion of electrophoresis, the proteins were transferred to an Immobilon P membrane (EMD Millipore, IPVH00010) at 25°C overnight. The next day, the membrane was blocked with 5% non-fat dry milk (BioRad, 180171A) in PBT (0.2% Tween 20 in PBS) for 1 hr at 25°C. Proteins were then labeled with the primary antibody of interest diluted in 5% milk in PBT at 25°C for 1 hr. After washing with PBT for 5 min three times, the membranes were incubated with secondary antibodies goat anti-rabbit IgG-HRP or goat anti-mouse IgG-HRP both diluted at 1:1000 in 5% milk in PBT for 1 hr at 25°C. After washing the membrane with PBT six times, the chemiluminescence signal was detected with a SuperSignal West Dura kit (Thermo Fisher Scientific, 34075) and X-ray films. The phospho-p38 antibody was used as follows to avoid the sequestration of the antibody by casein in the milk. The membrane was blocked with 5% non-fat dry milk in TBST (50 mM Tris-HCl pH 7.5, 0.1% Tween 20) for 1 hr at 25°C, followed by wash with TBST for 5 min three times. The p-p38 antibody was diluted in 5% BSA in TBST and three subsequent washes were done with TBST. Goat anti-rabbit IgG-HRP was diluted in 5% milk in TBST and six subsequent washes were done with TBST.

### Quantitative RT-PCR (qRT-PCR)

RNA was extracted from cells using a Quick RNA Microprep (Zymo Research, R1051) or RNeasy Plus Mini (Qiagen, 74136) kit, depending on cell number. RNA quantity and purity were assessed using a NanoDrop Lite spectrophotometer (Thermo Fisher Scientific). cDNA was synthesized with ProtoScript II Reverse Transcriptase (New England Biolabs, M0368L). Quantitative PCR (qPCR) was performed with the primers listed in Table S1 and a GoTaq qPCR Master Mix (Promega, A6002) in a Mastercycler realplex^2^ thermocycler (Eppendorf). PCR conditions were as follows: initial denaturation at 95°C for 10 min, 40 cycles of 95°C for 15 sec, 30 sec at the specific annealing temperature for each set of primers, and 72°C for 30 sec, and a melting curve step to check the specificity of the reaction. mRNA expression levels were analyzed by normalizing expression values to glyceraldehyde 3-phosphate dehydrogenase (*Gapdh*) expression. Mean ± SEM of biological triplicates with technical triplicates each were calculated.

Nascent mRNA was measured with a Click-iT Nascent RNA Capture kit (Thermo Fisher Scientific, C10365). C2C12 cells were incubated with 0.1 mM 5-ethynyl uridine (EU) for 4 hr, followed by the isolation of EU-labeled mRNAs with magnetic beads for cDNA synthesis and qPCR.

### ChIP-PCR

Two million C2C12 cells were treated with 1% paraformaldehyde for 10 min and then 125 mM glycine, followed by washing with PBS twice. Chromatin was prepared by the treatment with 300 μl cell lysis buffer (50 mM HEPES pH7.8, 85 mM NaCl, 0.5% NP-40, and cOmplete Mini Protease Inhibitor Cocktail (MilliporeSigma, 11 836 153 001)) for 15 min on ice with vortexing for 15 sec every 5 min. After centrifugation at 1,500 x *g* for 5 min at 4°C, the pellet was resuspended in 50 μl nuclear lysis buffer (50 mM Tris-HCl pH 8.0, 10 mM EDTA, 1% SDS, and cOmplete protease inhibitor cocktail) and incubated for 5 min on ice. Chromatin was sheared by sonication with a Bioruptor 300 (Diagenode) with 30 cycles of 30 sec-on and 30 sec-off with the high power setting at 4°C. After centrifugation at 15,000 x*g* for 15 min at 4°C, supernatant was incubated with 2 μg antibody, 2 μl Dynabeads Protein G (Thermo Fisher Scientific, 10004D), and 400 μl dilution buffer (20 mM Tris-HCl pH 8.0, 150 mM NaCl, 2 mM EDTA, 1% Triton X-100, and cOmplete protease inhibitor cocktail) for 16 hr at 4°C with rotation at 20 rpm. The beads were sequentially washed with 500 μl each of dilution buffer, LiCl buffer (20 mM Tris-HCl pH 8.0, 250 mM LiCl, 2 mM EDTA, 0.1% SDS, and 1% Triton X-100) twice, and TE (10 mM Tris-HCl and 1 mM EDTA) rotating for 5 min at 4°C each. The beads were resuspended in 100 μl elution solution (0.1 M NaHCO_3_, 1% SDS, and 200 μg/ml proteinase K) and incubated for 2 hr at 65°C with rotation to reverse crosslink. Proteinase K was inactivated by heating for 10 min at 95°C. DNA was purified with a ChIP DNA Clean & Concentrator (Zymo Research, D5205) and applied for qPCR as described above. Results of biological triplicates with technical duplicates each were presented as ratios in comparison to the results with normal mouse IgG as control.

The ChIP-seq data GSE25308 (Pol II, H3K4me1, H3K4me3, and H3K9ac), GSE37527 (H3K27ac and p300), and GSE108650 (Bmal1) were downloaded from the UCSC Mouse Genome Browser mm9. The quality of all downloaded data was evaluated using FastQC v0.11.5 (http://www.bioinformatics.babraham.ac.uk/projects/fastqc/) and adapter sequence was trimmed using Trimmomatic v0.33 (http://www.usadellab.org/cms/index.php?page=trimmomatic) (Bolger et al., 2014). The filtered high-quality reads were then mapped to a reference genome (GRCm38/mm10) using HISAT2 v2.0.2 (https://ccb.jhu.edu/software/hisat2/index.shtml) (Kim et al., 2015). Resulting BAM files with MACS v2.2.5 (https://github.com/macs3-project/MACS) were used to generate peaks (Zhang et al., 2008).

### 3C analysis

3C was performed combining two published protocols (Hagege et al., 2007; Naumova et al., 2012). To make a BAC control library, a mouse 110 kb BAC clone encoding *Nctc1* and *Igf2* was purchased from Thermo Fisher Scientific (RPCI23.C) and the DNA was prepared with a ZR BAC DNA Miniprep kit (Zymo Research (D4048)) as control. Ten micrograms of DNA was digested with BamHI for 16 hr, followed by phenol-chloroform extraction and ethanol precipitation twice. DNA fragments were ligated with T4 ligase for 16 hr at 16°C. DNA was purified with phenol-chloroform extraction and ethanol precipitation twice and used in qPCR.

C2C12 cells were treated with 1% paraformaldehyde and then with 125 mM glycine as described in the ChIP-PCR method. Ten million cells were treated with lysis buffer (10 mM Tris-HCl pH7.5, 10 mM NaCl, and 0.2% NP-40) for 10 min on ice. After centrifugation, the pellet was washed once with the digestion buffer for BamHI and incubated in a series of buffers for 1 hr at 37°C each: BamHI buffer, BamHI buffer with 0.3% SDS, and BamHI buffer with 0.3% SDS and 2% Triton X-100. Finally, the cells were incubated with 400 units BamHI for 16 hr at 37°C with rotation. BamHI was inactivated with 1.6 % SDS and incubation at 65°C for 30 min. The cell suspension was diluted 12-fold with lysis buffer (10 mM Tris-HCl pH7.5, 10 mM NaCl, 0.2% NP-4, and 1% Triton X-100) and incubated for 1 hr at 37°C with rotation. After the addition of 100 units T4 DNA ligase, the reaction mix was incubated for 4 hr at 16°C, followed by incubation for 1 hr at 25°C. Finally, the reaction mix was incubated for 16 hr at 65°C in the presence of 300 μg proteinase K, followed by phenol-chloroform extraction and ethanol precipitation twice as a 3C library.

PCR data was analyzed as detailed in reference (Naumova et al., 2012). Briefly, PCR was performed with 10 ng BAC control library and 3C library as templates using the 3C-0 kb primer in combination with each of 3C-76 kb through 3C-112 kb primers. The primer pair 3C-Clock1 and 3C-Clock2 at the *Clock* gene, which showed a consistent amplification throughout the circadian rhythms, was used as an internal control. PCR conditions were as it follows: initial denaturation at 95°C for 5 min, 35 cycles of 95°C for 30 sec, 70°C for 30 sec, and 72°C for 20 sec, and an extension at 72°C for 5 min. PCR products were resolved with a 2% agarose gel and stained with ethidium bromide. The images were captured with a Gel Logic 212 Pro system (CareStream Molecular Imaging) and the band intensity was quantified with Fiji. Biological triplicates were analyzed for each group.

### RNA-seq

Total RNA was prepared from KO C2C12 cells before differentiation (day 0) and day 3 and 5 during differentiation. RNA concentration and RNA Integrity Number (RIN) were measured with an Agilent BioAnalyzer 2100. Samples with RIN over 8 were used to create sequencing libraries at the University of Minnesota Genomics Center. One microgram of total RNA was used to create each sequencing library using a Truseq RNA Sample Preparation Kit (Illumina, RS-122-2001). Briefly, poly-adenylated RNA was first purified using oligo-dT-coated magnetic beads. RNA was then fragmented and reverse-transcribed into cDNA. The cDNA was further fragmented, blunt-ended, and ligated to barcoded adaptors and amplified. The final library size distribution was validated with capillary electrophoresis and quantified with a Quant-iT PicoGreen dsDNA Assay Kit (Thermo Fisher Scientific, P11496) and qPCR. Indexed libraries were pooled and size-selected to 320 bp ± 5% with a LabChip XT (PerkinElmer). Libraries were loaded onto a single-read flow cell and amplified on a cBot (Illumina) before sequencing using a NextSeq High (Illumina). The RNA-seq data have been deposited to Gene Expression Omnibus (GEO) under the accession number of GEO: GSE150785.

### Bioinformatics Analysis

On average, 58.78 million reads (51.23 million to 64.24 million) were generated per library. The demultiplexed FASTQ files were analyzed using a customized pipeline (gopher-pipelines; https://bitbucket.org/jgarbe/gopher-pipelines/overview) developed and maintained by the Minnesota Supercomputing Institute. Briefly, FastQC v0.11.5 (http://www.bioinformatics.babraham.ac.uk/projects/fastqc/) was used to check on the sequencing quality of the FASTQ files. Then adapters and low-quality reads were trimmed using Trimmomatic v0.33 (http://www.usadellab.org/cms/index.php?page=trimmomatic) (Bolger et al., 2014). An additional quality check with FastQC was performed on the post-trimming sequences to ensure successful adaptor and quality trimming. The remaining sequences were then aligned to the GRCm38/mm10 reference genome using HISAT2 v2.0.2 (https://ccb.jhu.edu/software/hisat2/index.shtml) and transcript abundance was counted using subread v1.4.6 (http://subread.sourceforge.net/) (Kim et al., 2015; Liao et al., 2014). Differential gene expression analysis was performed with R v3.6.2, *edgeR* (https://bioconductor.org/packages/release/bioc/html/edgeR.html) (Robinson et al., 2010). Gene ontology analysis of differentially expressed genes was performed by functionally annotate the genes and perform an overrepresentation enrichment test using PANTHER (http://pantherdb.org/) (Mi et al., 2013). Heat maps were generated using the log-transformed counts with *pheatmap* v1.0.12 (https://cran.r-project.org/web/packages/pheatmap/index.html) packages. Hierarchical clustering was performed using the average linkage clustering method with the correlation coefficient as a similarity metric.

### Principal component analysis

The principal component analysis was performed to investigate the clustering of the datasets. The transformed and normalized gene expression values were used and principal components were computed with R v3.6.2., prcomp() (https://www.rdocumentation.org/packages/utils/versions/3.6.2/topics/prompt). The first and second largest variance components in the data (PC1 and PC2) were visualized as a scatter plot using R v3.6.2, ggplot2 v3.2.1 (https://cran.r-project.org/web/packages/ggplot2/index.html).

### Scattered plot

Regression analysis was used to evaluate correlations between the two groups. Normalized expression values of each group were transformed with natural log for better linear fitting and plotted against each other. R-squared value was calculated and shown on each regression line. All plots were produced in R v3.6.2, ggplot2 v3.2.1.

### Statistical analyses

One-sided Student’s t-tests were used in the analysis of statistical significance of the difference in the differentiation index, the fusion index, EdU uptake, qRT-PCR, and ChIP-PCR data. The mean + or ± SEM obtained from biological triplicates was shown in each graph unless stated otherwise. 24 hr rhythmicity was detected with Cosinor at DiscoRhythm (https://mcarlucci.shinyapps.io/discorhythm/) (Carlucci et al., 2019).

## Acknowledgements

We thank James Staats, Brian Ruis, and Allison Keith for technical supports, Toshio Kitamura for the pMXs-IP plasmid and the PLAT-E cells, and Karyn A Esser for critical reading of the manuscript. We acknowledge the Minnesota Supercomputing Institute, University of Minnesota Informatics Institute, and University of Minnesota Genomics Center for providing high-performance computing resources and the gopher-pipelines. S.L. and C.Y. were supported by the Minnesota Stem Cell Institute. A.A. was supported by the NIH (R01AR062142 and R21AR070319). N.K was supported by the NIH (R01GM137603 and R21AR076167), Regenerative Medicine Minnesota (RMM 101617 DS 004), and Grant-in-Aid of Research University of Minnesota (291987). The content is solely the responsibility of the authors and does not necessarily represent the official views of the NIH.

## Author contribution

N. Katoku-Kikyo, E. Paatela, D.L. Houtz, B. Lee, D. Munson, X. Wang, M. Hussein, J. Bhatia, and Y. Asakura: investigation. S. Lim and C. Yuan: formal analysis. A. Asakura and N. Kikyo: conceptualization, funding acquisition, investigation, project administration, supervision, and writing of the original draft. All: review and editing of the manuscript.

## Supplemental Figure Legends

**Figure S1.**
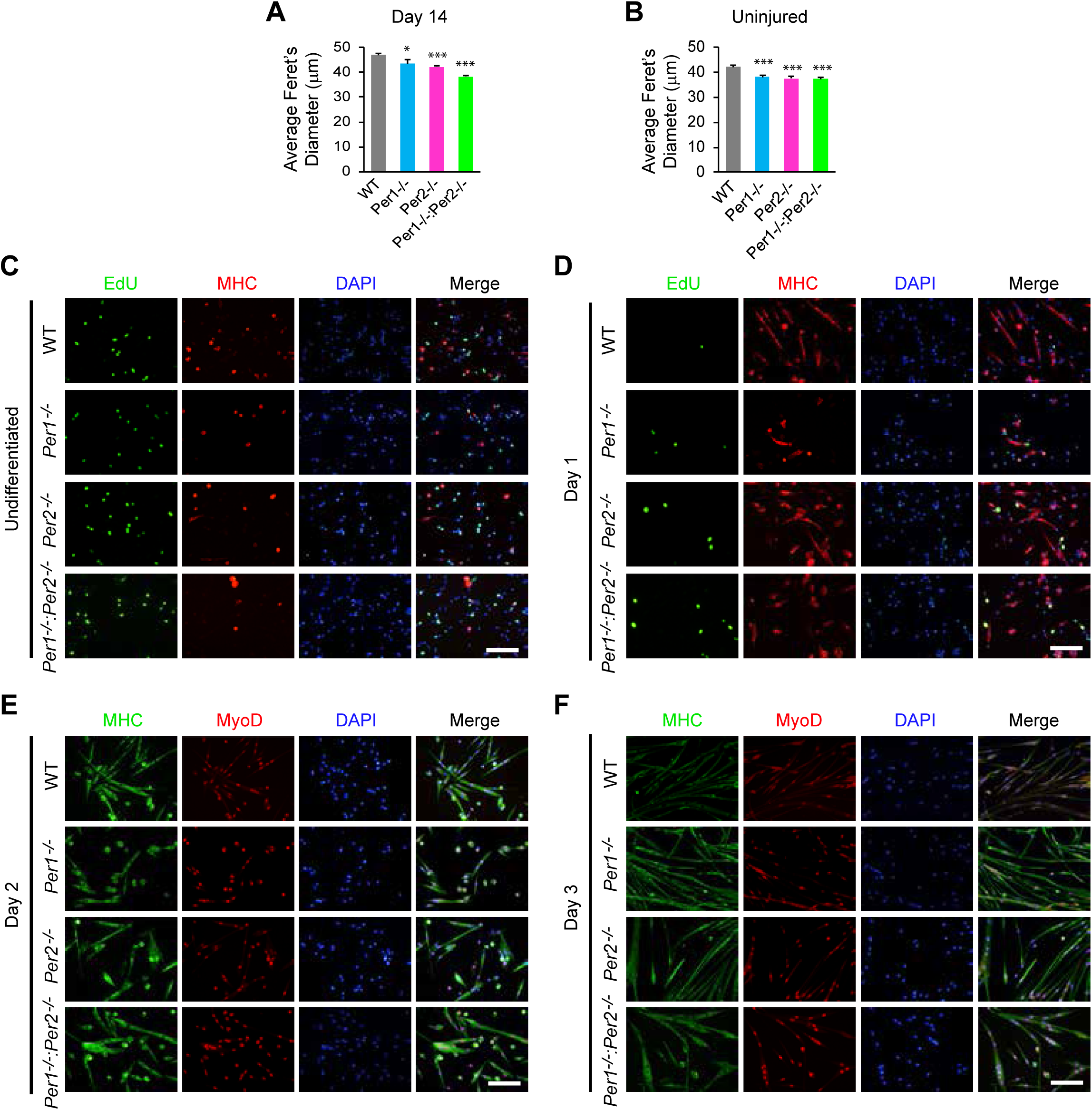
Regeneration of TA muscle and myoblast differentiation comparing *Per1^-/-^*, *Per2^- /-^*, and *Per1^-/-^:Per2^-/-^* mice. (A) Average of the minimal Feret’s diameters of myofibers with centrally-located nuclei on day 14. TA muscle was injured with barium chloride at ZT14 on day 0 and harvested 14 days later. n = 8 mice with 4 males and 4 females in each group in (A) and (B). (B) Average of the minimal Feret’s diameters of myofibers in uninjured mice. (C)-(F) EdU uptake and immunofluorescence staining of undifferentiated (C) and differentiating primary myoblasts on day 1 (D), day 2 (E), and day 3 (F) with antibodies against MHC and MyoD. DNA was counterstained with DAPI. Bar, 100 μm. * p < 0.05 and *** p < 0.001 with Student’s t-test in comparison to WT mice. Data are presented as mean + SEM.

**Figure S2.**
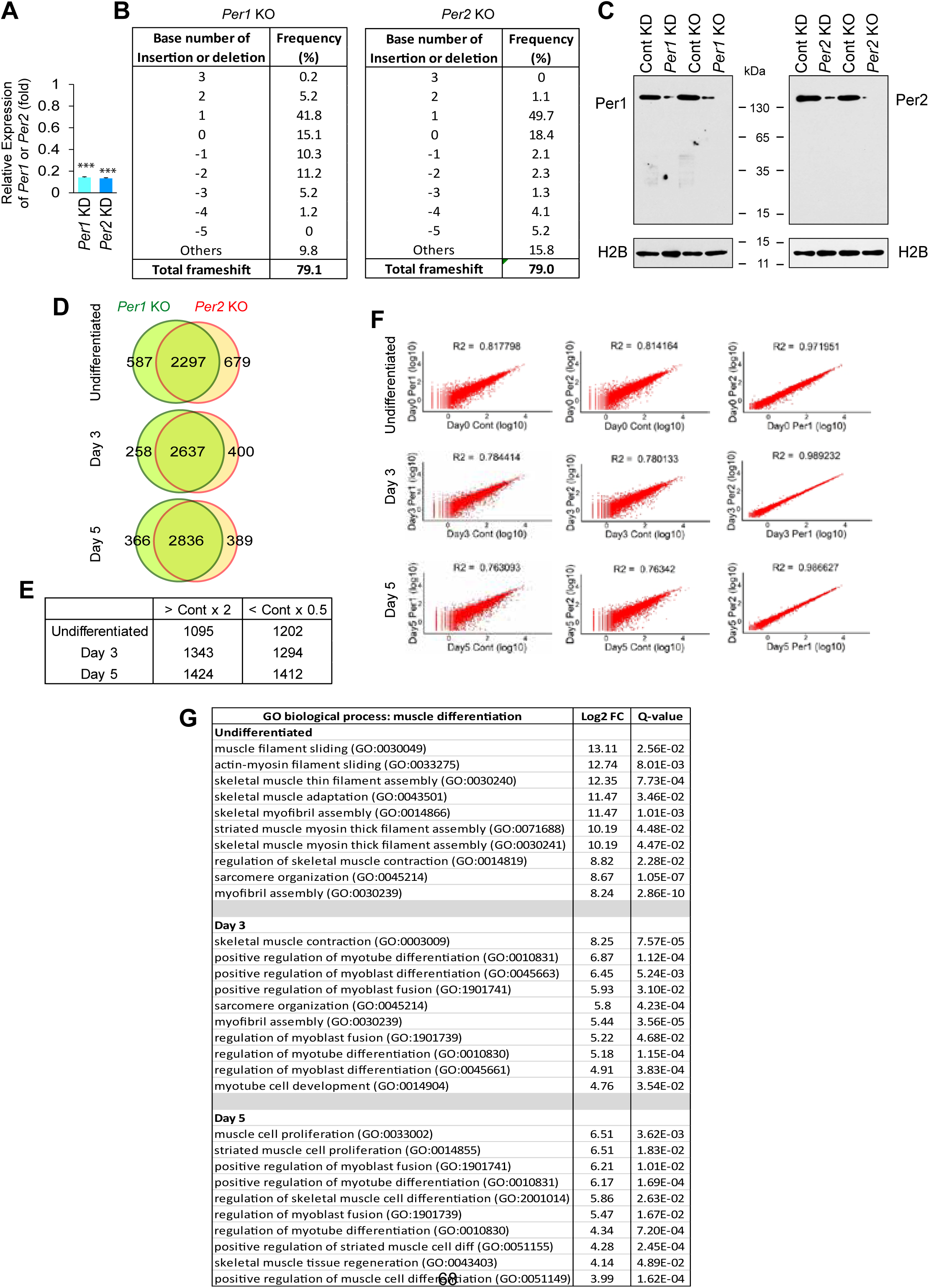
Comparison of transcriptomes between control and Per-depleted C2C12 cells. (A) Relative expression levels of *Per1* and *Per2* mRNAs in C2C12 cells after KD of each gene. The expression level with control scrambled shRNA was defined as 1.0 for each gene. *** p<0.001 with Student’s t-test. Data are presented as mean + SEM of biological triplicates. (B) Indel frequency in *Per1* KO and *Per2* KO C2C12 cells analyzed with the TIDE software (https://tide.nki.nl/). (C) Western blotting demonstrating downregulation of Per1 and Per2 in KD and KO cells. Histone H2B was used as loading control. (D) Venn diagrams displaying the number of genes whose expression levels were >200% or <50% of those of control KO cells. n = 1 in (D)-(G). (E) The number of genes that were commonly up- (> Cont x 2) or down-regulated (< Cont x 0.5) more than twofold in *Per1* KO and *Per2* KO cells compared with control KO cells. (F) Scatter plots comparing control, *Per1* KO, and *Per2* KO C2C12 cells. (G) Gene ontology (GO) terms relevant to muscle differentiation that were enriched in the genes commonly downregulated in *Per1* KO and *Per2* KO cells compared with control KO cells.

**Figure S3.**
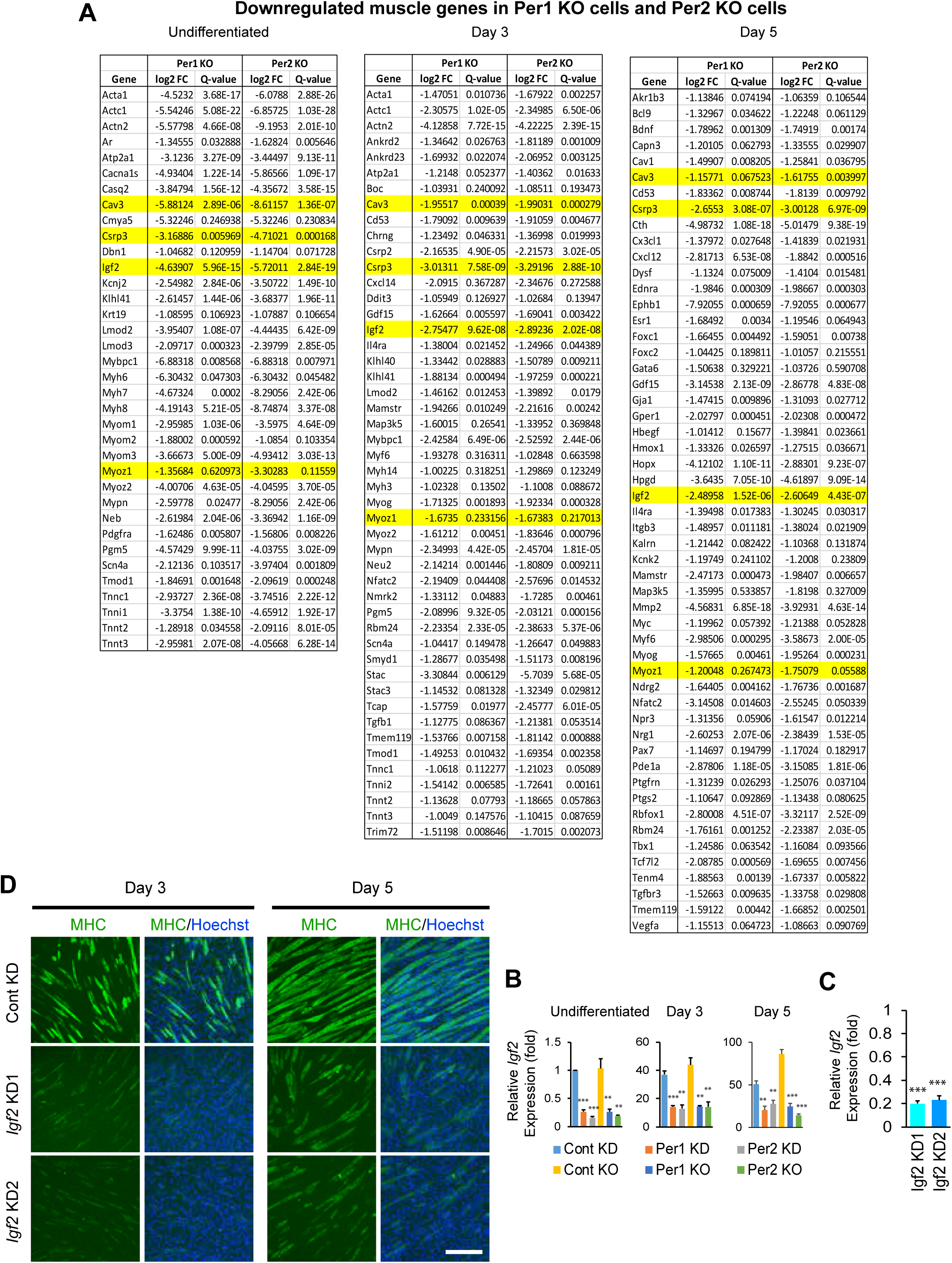
RNA-seq analysis of *Per1* and *Per2* KO cells and differentiation of *Igf2* KD cells. (A) List of genes belonging to the GO terms shown in Figure S2G. Four genes commonly downregulated in *Per1* KO and *Per2* KO cells in an undifferentiated state and during differentiation are highlighted in yellow. n = 1 in (A) and (B). (B) Relative expression level of *Igf2* mRNA determined by qPCR in C2C12 cells after depletion of *Per1* and *Per2*. The value obtained with control scrambled shRNA on day 0 (before differentiation) was defined as 1.0. (C) Relative expression level of *Igf2* mRNA after KD with two shRNA clones. The expression level with control shRNA was defined as 1.0. (D) MHC staining of differentiating C2C12 cells after *Igf2* KD with two shRNAs. Bar, 200 μm. ** p < 0.01 and *** p < 0.001 with Student’s t-test in comparison to control cells. Data are presented as mean + SEM of biological triplicates.

**Figure S4.**
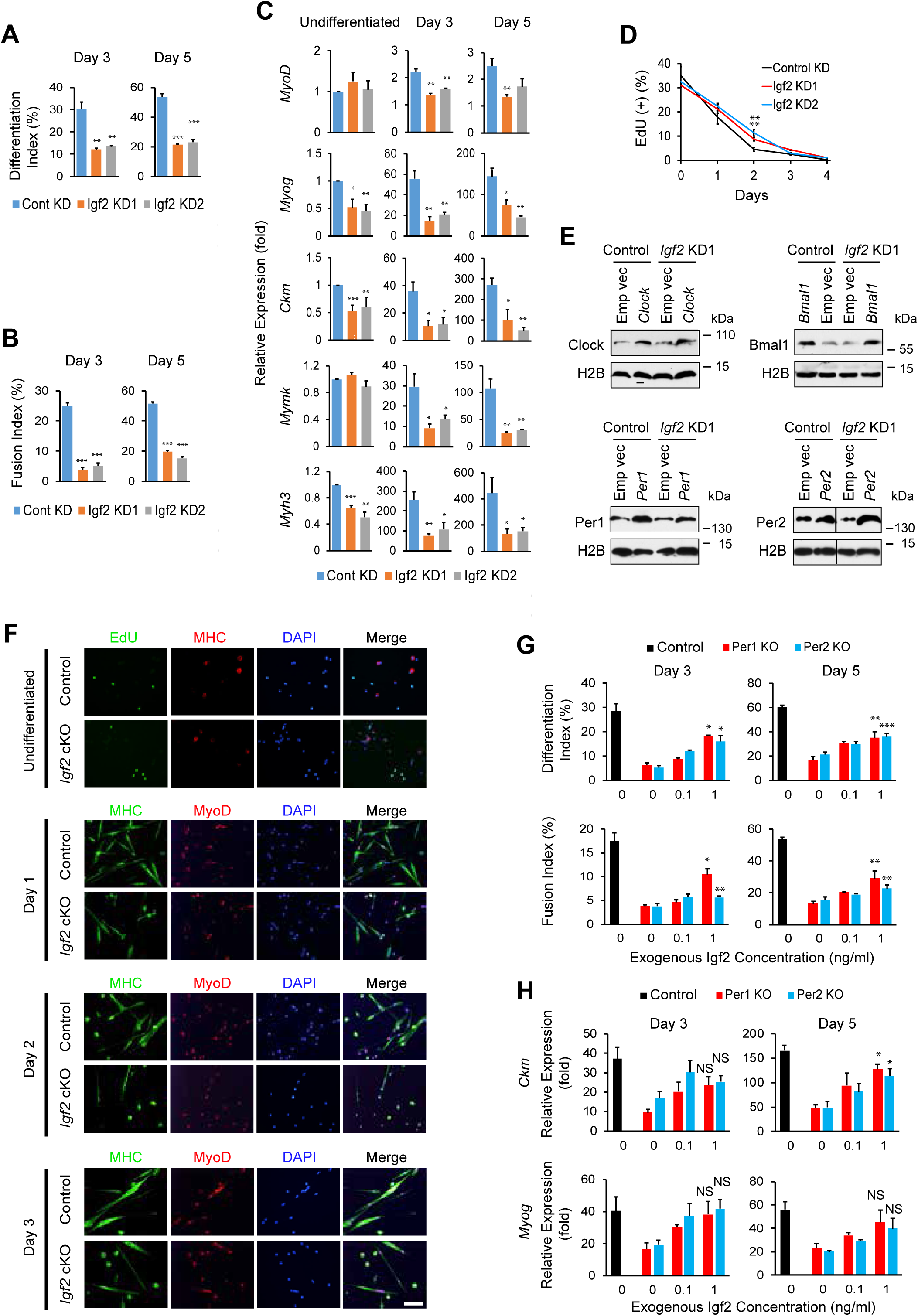
Per1/Per2-Igf2 axis and myoblast differentiation. (A) and (B) Differentiation index (A) and fusion index (B) during differentiation of *Igf2* KD cells. (C) Relative expression levels of five muscle genes during differentiation of *Igf2* KD cells. The value obtained with day 0 control KD cells was defined as 1.0. (D) Temporal profile of the frequency of EdU(+) nuclei in *Igf2* KD cells during differentiation. (E) Western blotting of Clock, Bmal1, Per1, and Per2 after retrovirus-mediated transduction of these genes in C2C12 cells. Empty vector (Emp Vec) was used as control. Cells transduced with *Igf2* shRNA clone 1 (Igf2 KD1) and those with control shRNA were compared. Histone H2B was used as loading control. (F) EdU uptake and immunofluorescence staining of undifferentiated and differentiating primary myoblasts with antibodies against MHC and MyoD. DNA was counterstained with DAPI. Bar, 100 μm. (G) Differentiation index (top) and fusion index (bottom) of control and *Per* KO C2C12 cells cultured with exogenous Igf2 at different concentrations for 3 and 5 days. Igf2 was not added to the control cells. (H) Relative expression levels of *Ckm* (top) and *Myog* (bottom) of control and *Per* KO C2C12 cells treated with exogenous Igf2 for 3 and 5 days. The expression level of the control cells before differentiation was defined as 1.0. Data are presented as mean + or ± SEM of biological triplicates. Each replicate includes n = 1,000-1,500 nuclei in (A) and (B). * p < 0.05, ** p < 0.01, and *** p < 0.001 with Student’s t-test in comparison to control. NS indicates that there was no statistically significant difference (p ≥ 0.05). Asterisks were only added to the values with 1 ng/ml Igf2 in (G) and (H).

**Figure S5.**
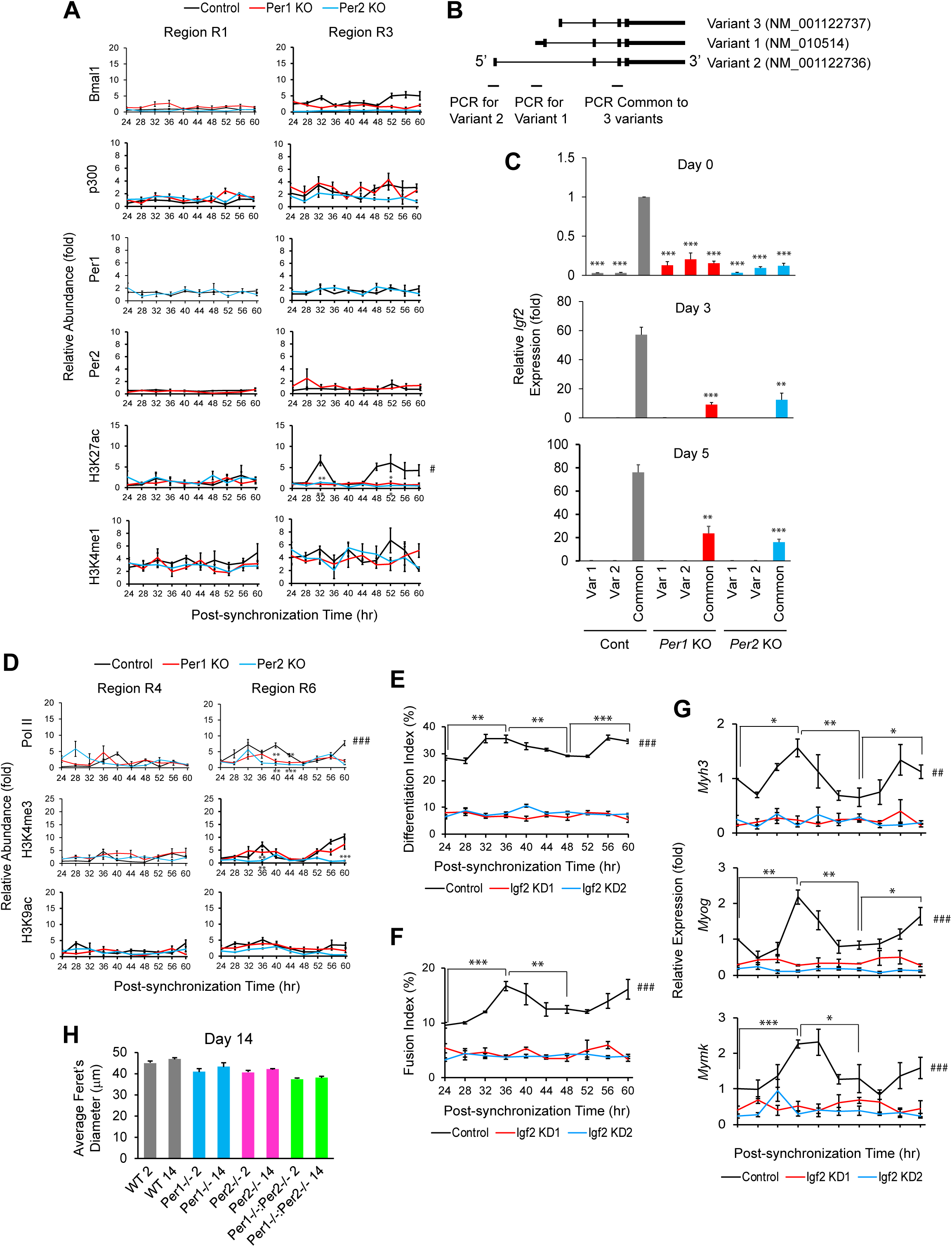
Per1/Per2-Igf2 axis and circadian timing-dependent myogenesis and muscle regeneration. (A) ChIP-PCR analyses of indicated proteins in control and *Per* KO C2C12 cells at the regions R1 and R3 shown in Fig. 5 F. Relative abundance compared with control IgG is shown. Peak time points when the control cell values were higher than those of *Per1* KO and *Per2* KO cells are highlighted with asterisks. (B) Locations of the PCR primers specific to two variants and common to all three variants of *Igf2*. (C) qPCR results of the *Igf2* variant mRNAs in control and *Per* KO cells. The PCR products obtained with the common primers largely represented the expression levels of variant 3 because the levels of variants 1 and 2 were by far lower than the level of variant 3. (D) ChIP-PCR analyses of indicated proteins in control and *Per* KO C2C12 cells at the regions R4 and R6 in Fig. 5 H. Relative abundance compared with control IgG is shown. Peak time points when the control cell values were higher than those of *Per1* KO and *Per2* KO cells are highlighted with asterisks. (E)-(G) Analyses of differentiation index (E), fusion index (F), and relative expression levels of differentiation-specific genes (G) with C2C12 cells that were induced to differentiate at the indicated post-synchronization time points. Control and *Igf2* KD cells prepared with two shRNA clones were compared. (H) Average Feret’s diameters of myofibers with centrally-located nuclei on day 14. TA muscle was injured with barium chloride at ZT2 or ZT14 on day 0 and harvested 14 days later. Mean + SEM of 8 mice including 4 males and 4 females in each group is shown. 2 and 14 at the end of each genotype indicate the injury time at ZT2 and ZT14, respectively. Data are presented as mean + or ± SEM of biological triplicates. Each replicate includes n = 1,000-1,500 nuclei in (E) and (F). * p < 0.05, ** p < 0.01, and *** p < 0.001 with Student’s t-test compared with control values in (A) and (D), control common primers in (B), and different time points in control cells in (E)-(G). ^#^ 24 h rhythmicity with Cosinor (p < 0.05), ^##^ P < 0.01, and ^###^ P < 0.001.

**Table S1.**
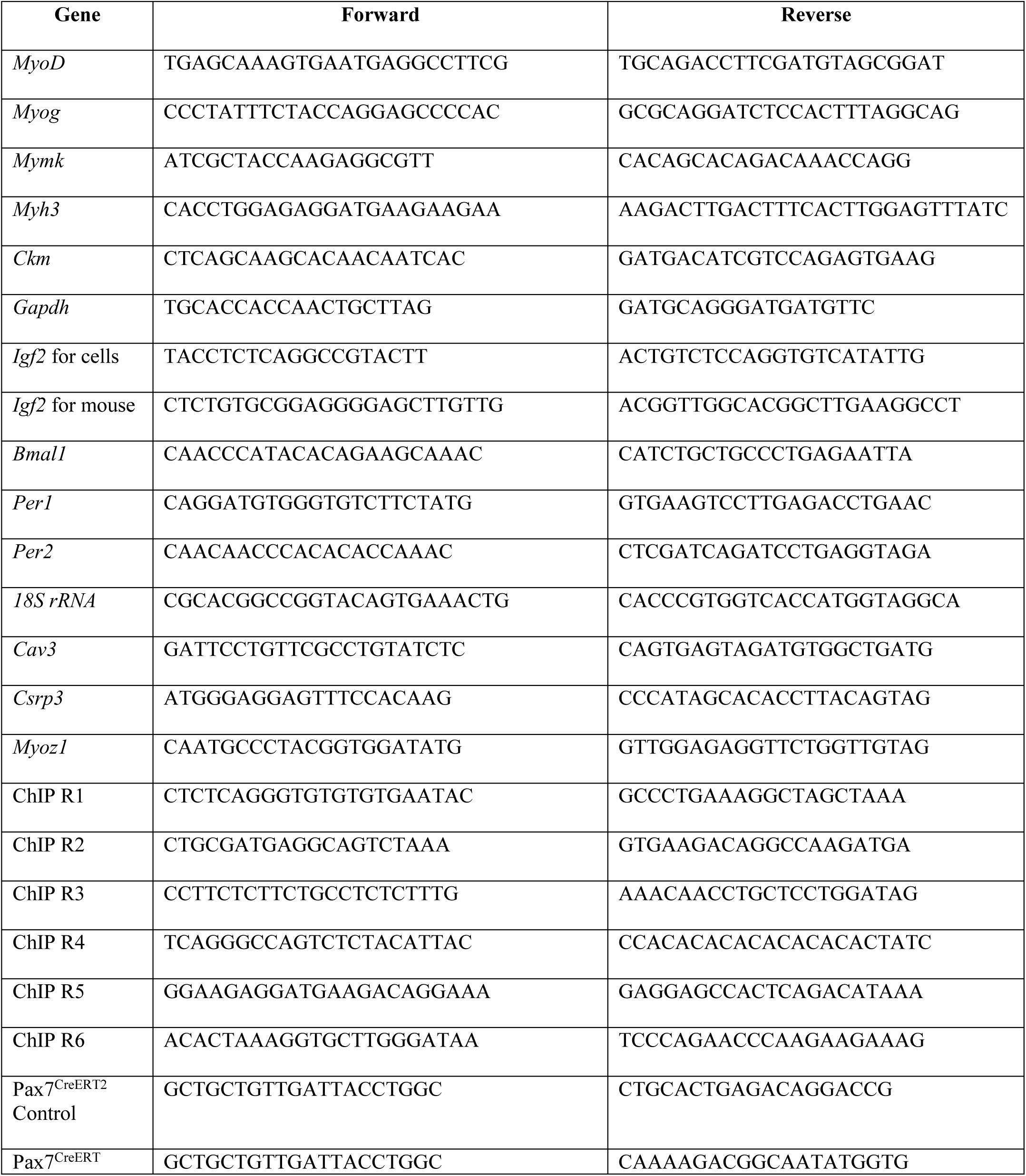
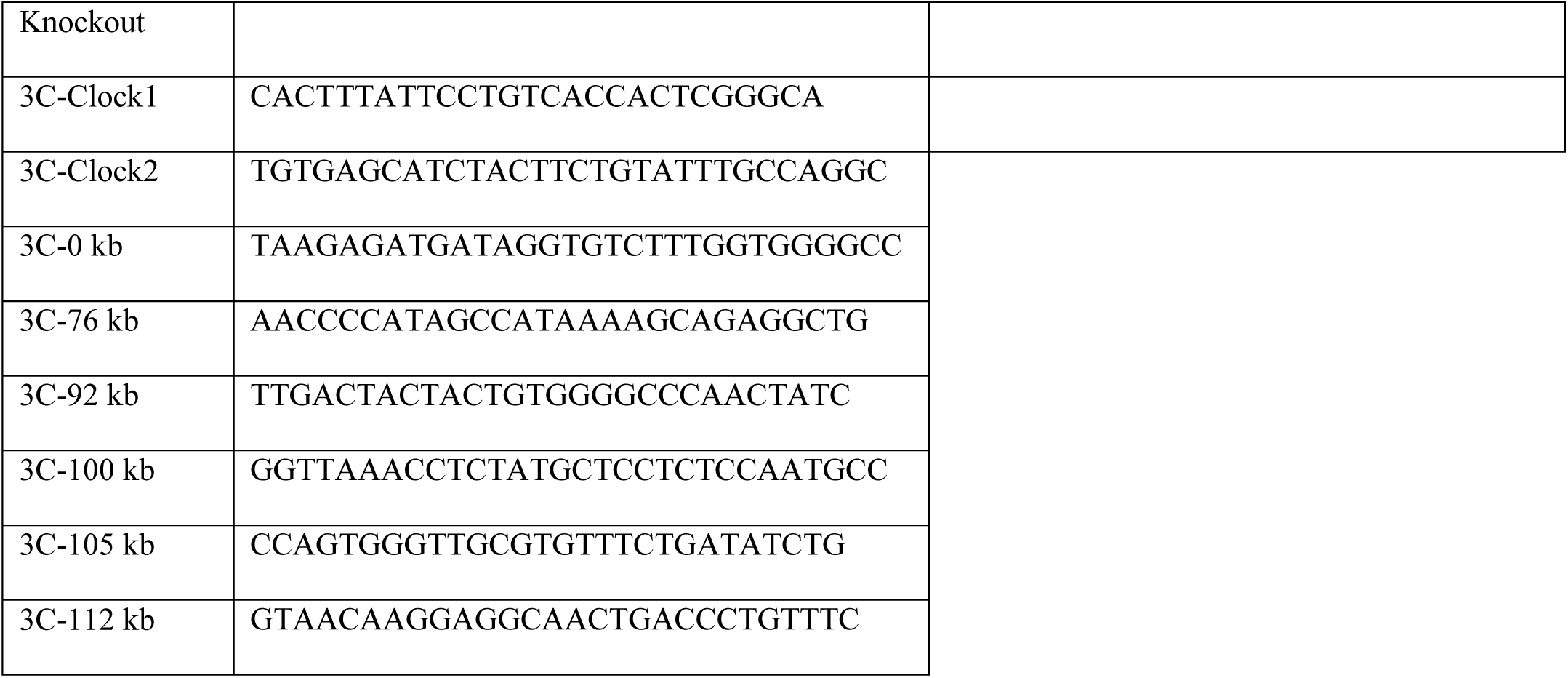
Sequences of PCR primers.

**Table S2.** Antibodies.

| **Protein** | **Manufacturer** | **Catalog #** | **Dilution (fold)** |
| --- | --- | --- | --- |
| CD31-PE | eBioscience | 12-0311 | 200 |
| CD45-PE | eBioscience | 12-0451 | 200 |
| Sca1-PE | eBioscience | 12-5981 | 200 |
| Integrin $\alpha 7$-biotin | Miltenyi Biotec | 130-102-125 | 200 |
| Laminin | MilliporeSigma | L0663 | 1000 |
| Bmal1 | Abcam | Ab3350 | 200 |
| Per1 | MilliporeSigma | AB2201 | 100 |
| Per2 | NOVUS | NB100-125 | 100 |
| Igf2 | Santa Cruz<br>Biotechnology | sc-5622 | 100 |
| p38$\alpha/\beta$ | Santa Cruz<br>Biotechnology | sc-7972 | 300 |
| p-p38 | Thermo Fisher Scientific | MA5-15177 | 400 |
| Histone H2B | Thermo Fisher Scientific | MA5-14835 | 1000 |
| H3K27ac | MilliporeSigma | 07-360 | 500 |
| H3K4me1 | MilliporeSigma | 07-436 | 500 |
| H3K4me3 | MilliporeSigma | 07-473 | 500 |
| H3K9ac | MilliporeSigma | 07-352 | 400 |
| p300 | MilliporeSigma | 05-257 | 300 |
| Pol II, CTD | MilliporeSigma | 05-952-I-100UG | 600 |
| Embryonic MHC | Developmental Studies<br>Hybridoma Bank | F1.652 | 50 |
| MHC | Developmental Studies<br>Hybridoma Bank | MF 20 | 50 |
| MyoD | Santa Cruz<br>Biotechnology | sc-304 | 500 |
| Alexa Fluor 488 goat anti-mouse IgG | Thermo Fisher Scientific | A-11029 | 1000 |
| Alexa Fluor 488 donkey anti-mouse IgG | Thermo Fisher Scientific | A-21202 | 1000 |
| Alexa Fluor 594 donkey anti-rat IgG | Thermo Fisher Scientific | A-21209 | 1000 |
| Alexa Fluor 594 donkey anti-rabbit IgG | Thermo Fisher Scientific | A-21207 | 1000 |
| Goat anti-mouse IgG, HRP labeled | Santa Cruz<br>Biotechnology | sc-2005 | 1000 |
| Goat anti-rabbit IgG, HRP labeled | Santa Cruz<br>Biotechnology | sc-2004 | 1000 |
| Normal mouse IgG | Santa Cruz<br>Biotechnology | sc-2025 | N/A |

**Table S3.**
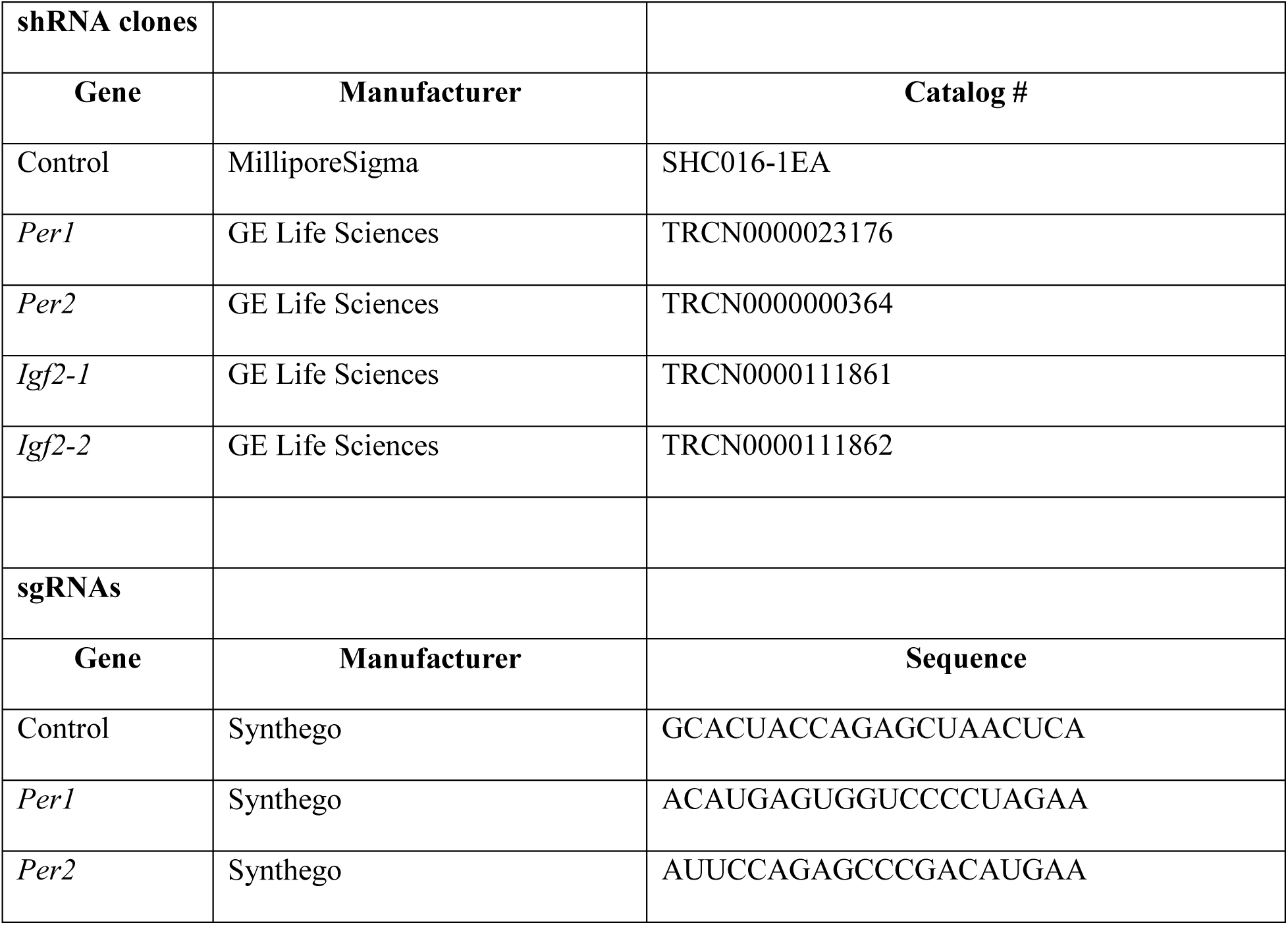
shRNA and sgRNA sequences.

## References

Akashi, M., A. Okamoto, Y. Tsuchiya, T. Todo, E. Nishida, and K. Node. 2014. A positive role for PERIOD in mammalian circadian gene expression. Cell Rep. 7:1056–1064.http://www.ncbi.nlm.nih.gov/pubmed/24794436

Allard, J.B., and C. Duan. 2018. IGF-Binding Proteins: Why Do They Exist and Why Are There So Many? Front Endocrinol (Lausanne). 9:117.http://www.ncbi.nlm.nih.gov/pubmed/29686648

Alzhanov, D., and P. Rotwein. 2016. Characterizing a distal muscle enhancer in the mouse Igf2 locus. Physiol Genomics. 48:167–172.http://www.ncbi.nlm.nih.gov/pubmed/26645089

Alzhanov, D.T., S.F. McInerney, and P. Rotwein. 2010. Long range interactions regulate Igf2 gene transcription during skeletal muscle differentiation. J Biol Chem. 285:38969–38977.http://www.ncbi.nlm.nih.gov/pubmed/20937833

Andrews, J.L., X. Zhang, J.J. McCarthy, E.L. McDearmon, T.A. Hornberger, B. Russell, K.S. Campbell, S. Arbogast, M.B. Reid, J.R. Walker, J.B. Hogenesch, J.S. Takahashi, and K.A. Esser. 2010. CLOCK and BMAL1 regulate MyoD and are necessary for maintenance of skeletal muscle phenotype and function. Proc Natl Acad Sci U S A. 107:19090–19095.http://www.ncbi.nlm.nih.gov/pubmed/20956306

Aoyama, S., S. Kojima, K. Sasaki, R. Ishikawa, M. Tanaka, T. Shimoda, Y. Hattori, N. Aoki, K. Takahashi, R. Hirooka, M. Takizawa, A. Haraguchi, and S. Shibata. 2018. Day-Night Oscillation of Atrogin1 and Timing-Dependent Preventive Effect of Weight-Bearing on Muscle Atrophy. EBioMedicine. 37:499–508.http://www.ncbi.nlm.nih.gov/pubmed/30391495

Aoyama, S., and S. Shibata. 2017. The Role of Circadian Rhythms in Muscular and Osseous Physiology and Their Regulation by Nutrition and Exercise. Front Neurosci. 11:63.https://www.ncbi.nlm.nih.gov/pubmed/28261043

Aoyama, S., and S. Shibata. 2020. Time-of-Day-Dependent Physiological Responses to Meal and Exercise. Front Nutr. 7:18.http://www.ncbi.nlm.nih.gov/pubmed/32181258

Asakura, A., P. Seale, A. Girgis-Gabardo, and M.A. Rudnicki. 2002. Myogenic specification of side population cells in skeletal muscle. J Cell Biol. 159:123–134.http://www.ncbi.nlm.nih.gov/entrez/query.fcgi?cmd=Retrieve&db=PubMed&dopt=Citation&list_uids=12379804

Bader, D., T. Masaki, and D.A. Fischman. 1982. Immunochemical analysis of myosin heavy chain during avian myogenesis in vivo and in vitro. J Cell Biol. 95:763–770.http://www.ncbi.nlm.nih.gov/pubmed/6185504

Bae, K., X. Jin, E.S. Maywood, M.H. Hastings, S.M. Reppert, and D.R. Weaver. 2001. Differential functions of mPer1, mPer2, and mPer3 in the SCN circadian clock. Neuron. 30:525–536.https://www.ncbi.nlm.nih.gov/pubmed/11395012

Bae, K., K. Lee, Y. Seo, H. Lee, D. Kim, and I. Choi. 2006. Differential effects of two period genes on the physiology and proteomic profiles of mouse anterior tibialis muscles. Mol Cells. 22:275–284.http://www.ncbi.nlm.nih.gov/pubmed/17202855

Baker, J., J.P. Liu, E.J. Robertson, and A. Efstratiadis. 1993. Role of insulin-like growth factors in embryonic and postnatal growth. Cell. 75:73–82.http://www.ncbi.nlm.nih.gov/pubmed/8402902

Baral, K., and P. Rotwein. 2019. The insulin-like growth factor 2 gene in mammals: Organizational complexity within a conserved locus. PLoS One. 14:e0219155.http://www.ncbi.nlm.nih.gov/pubmed/31251794

Baumert, P., M.J. Lake, C.E. Stewart, B. Drust, and R.M. Erskine. 2016. Genetic variation and exercise-induced muscle damage: implications for athletic performance, injury and ageing. Eur J Appl Physiol. 116:1595–1625.http://www.ncbi.nlm.nih.gov/pubmed/27294501

Benitah, S.A., and P.S. Welz. 2020. Circadian Regulation of Adult Stem Cell Homeostasis and Aging. Cell Stem Cell. 26:817–831.https://www.ncbi.nlm.nih.gov/pubmed/32502402

Bolger, A.M., M. Lohse, and B. Usadel. 2014. Trimmomatic: a flexible trimmer for Illumina sequence data. Bioinformatics. 30:2114–2120.https://www.ncbi.nlm.nih.gov/pubmed/24695404

Bose, S., and F.R. Boockfor. 2010. Episodes of prolactin gene expression in GH3 cells are dependent on selective promoter binding of multiple circadian elements. Endocrinology. 151:2287–2296.http://www.ncbi.nlm.nih.gov/pubmed/20215567

Carlucci, M., A. Krisciunas, H. Li, P. Gibas, K. Koncevicius, A. Petronis, and G. Oh. 2019. DiscoRhythm: an easy-to-use web application and R package for discovering rhythmicity. Bioinformatics.https://www.ncbi.nlm.nih.gov/pubmed/31702788

Cermakian, N., L. Monaco, M.P. Pando, A. Dierich, and P. Sassone-Corsi. 2001. Altered behavioral rhythms and clock gene expression in mice with a targeted mutation in the Period1 gene. EMBO J. 20:3967–3974.http://www.ncbi.nlm.nih.gov/pubmed/11483500

Chatterjee, S., D. Nam, B. Guo, J.M. Kim, G.E. Winnier, J. Lee, R. Berdeaux, V.K. Yechoor, and K. Ma. 2013. Brain and muscle Arnt-like 1 is a key regulator of myogenesis. J Cell Sci. 126:2213–2224.https://www.ncbi.nlm.nih.gov/pubmed/23525013

Chatterjee, S., H. Yin, W. Li, J. Lee, V.K. Yechoor, and K. Ma. 2019. The Nuclear Receptor and Clock Repressor Rev-erbalpha Suppresses Myogenesis. Sci Rep. 9:4585.https://www.ncbi.nlm.nih.gov/pubmed/30872796

Chatterjee, S., H. Yin, D. Nam, Y. Li, and K. Ma. 2015. Brain and muscle Arnt-like 1 promotes skeletal muscle regeneration through satellite cell expansion. Exp Cell Res. 331:200–210.https://www.ncbi.nlm.nih.gov/pubmed/25218946

Chiou, Y.Y., Y. Yang, N. Rashid, R. Ye, C.P. Selby, and A. Sancar. 2016. Mammalian Period represses and de-represses transcription by displacing CLOCK-BMAL1 from promoters in a Cryptochrome-dependent manner. Proc Natl Acad Sci U S A. 113:E6072–E6079.http://www.ncbi.nlm.nih.gov/pubmed/27688755

Creyghton, M.P., A.W. Cheng, G.G. Welstead, T. Kooistra, B.W. Carey, E.J. Steine, J. Hanna, M.A. Lodato, G.M. Frampton, P.A. Sharp, L.A. Boyer, R.A. Young, and R. Jaenisch. 2010. Histone H3K27ac separates active from poised enhancers and predicts developmental state. Proc Natl Acad Sci U S A. 107:21931–21936.http://www.ncbi.nlm.nih.gov/pubmed/21106759

Crosby, P., R. Hamnett, M. Putker, N.P. Hoyle, M. Reed, C.J. Karam, E.S. Maywood, A. Stangherlin, J.E. Chesham, E.A. Hayter, L. Rosenbrier-Ribeiro, P. Newham, H. Clevers, D.A. Bechtold, and J.S. O’Neill. 2019. Insulin/IGF-1 Drives PERIOD Synthesis to Entrain Circadian Rhythms with Feeding Time. Cell. 177:896–909 e820.http://www.ncbi.nlm.nih.gov/pubmed/31030999

D’Alessandro, M., S. Beesley, J.K. Kim, R. Chen, E. Abich, W. Cheng, P. Yi, J.S. Takahashi, and C. Lee. 2015. A tunable artificial circadian clock in clock-defective mice. Nat Commun. 6:8587.http://www.ncbi.nlm.nih.gov/pubmed/26617050

Devaney, J.M., E.P. Hoffman, H. Gordish-Dressman, A. Kearns, E. Zambraski, and P.M. Clarkson. 2007. IGF-II gene region polymorphisms related to exertional muscle damage. J Appl Physiol (1985). 102:1815–1823.http://www.ncbi.nlm.nih.gov/pubmed/17289909

Dierickx, P., L.W. Van Laake, and N. Geijsen. 2018. Circadian clocks: from stem cells to tissue homeostasis and regeneration. EMBO reports. 19:18–28.http://www.ncbi.nlm.nih.gov/pubmed/29258993 http://www.pubmedcentral.nih.gov/articlerender.fcgi?artid=PMC5757216

Duan, C., H. Ren, and S. Gao. 2010. Insulin-like growth factors (IGFs), IGF receptors, and IGF-binding proteins: roles in skeletal muscle growth and differentiation. Gen Comp Endocrinol. 167:344–351.http://www.ncbi.nlm.nih.gov/pubmed/20403355

Dyar, K.A., M.J. Hubert, A.A. Mir, S. Ciciliot, D. Lutter, F. Greulich, F. Quagliarini, M. Kleinert, K. Fischer, T.O. Eichmann, L.E. Wright, M.I. Pena Paz, A. Casarin, V. Pertegato, V. Romanello, M. Albiero, S. Mazzucco, R. Rizzuto, L. Salviati, G. Biolo, B. Blaauw, S. Schiaffino, and N.H. Uhlenhaut. 2018. Transcriptional programming of lipid and amino acid metabolism by the skeletal muscle circadian clock. PLoS Biol. 16:e2005886.http://www.ncbi.nlm.nih.gov/pubmed/30096135

Florini, J.R., K.A. Magri, D.Z. Ewton, P.L. James, K. Grindstaff, and P.S. Rotwein. 1991. “Spontaneous” differentiation of skeletal myoblasts is dependent upon autocrine secretion of insulin-like growth factor-II. J Biol Chem. 266:15917–15923.http://www.ncbi.nlm.nih.gov/pubmed/1651927

Gardner, S., S.M. Gross, L.L. David, J.E. Klimek, and P. Rotwein. 2015. Separating myoblast differentiation from muscle cell fusion using IGF-I and the p38 MAP kinase inhibitor SB202190. Am J Physiol Cell Physiol. 309:C491–500.http://www.ncbi.nlm.nih.gov/pubmed/26246429

Gumz, M.L., L.R. Stow, I.J. Lynch, M.M. Greenlee, A. Rudin, B.D. Cain, D.R. Weaver, and C.S. Wingo. 2009. The circadian clock protein Period 1 regulates expression of the renal epithelial sodium channel in mice. J Clin Invest. 119:2423–2434.http://www.ncbi.nlm.nih.gov/pubmed/19587447

Gustafson, C.L., and C.L. Partch. 2015. Emerging models for the molecular basis of mammalian circadian timing. Biochemistry. 54:134–149.https://www.ncbi.nlm.nih.gov/pubmed/25303119

Hagege, H., P. Klous, C. Braem, E. Splinter, J. Dekker, G. Cathala, W. de Laat, and T. Forne. 2007. Quantitative analysis of chromosome conformation capture assays (3C-qPCR). Nat Protoc. 2:1722–1733.http://www.ncbi.nlm.nih.gov/pubmed/17641637

Harfmann, B.D., E.A. Schroder, and K.A. Esser. 2015. Circadian rhythms, the molecular clock, and skeletal muscle. J Biol Rhythms. 30:84–94.http://www.ncbi.nlm.nih.gov/pubmed/25512305

Hirano, A., Y.H. Fu, and L.J. Ptacek. 2016. The intricate dance of post-translational modifications in the rhythm of life. Nat Struct Mol Biol. 23:1053–1060.https://www.ncbi.nlm.nih.gov/pubmed/27922612

Hodge, B.A., Y. Wen, L.A. Riley, X. Zhang, J.H. England, B.D. Harfmann, E.A. Schroder, and K.A. Esser. 2015. The endogenous molecular clock orchestrates the temporal separation of substrate metabolism in skeletal muscle. Skelet Muscle. 5:17.http://www.ncbi.nlm.nih.gov/pubmed/26000164

Hodge, B.A., X. Zhang, M.A. Gutierrez-Monreal, Y. Cao, D.W. Hammers, Z. Yao, C.A. Wolff, P. Du, D. Kemler, A.R. Judge, and K.A. Esser. 2019. MYOD1 functions as a clock amplifier as well as a critical co-factor for downstream circadian gene expression in muscle. Elife. 8.http://www.ncbi.nlm.nih.gov/pubmed/30789342

Hoyle, N.P., E. Seinkmane, M. Putker, K.A. Feeney, T.P. Krogager, J.E. Chesham, L.K. Bray, J.M. Thomas, K. Dunn, J. Blaikley, and J.S. O’Neill. 2017. Circadian actin dynamics drive rhythmic fibroblast mobilization during wound healing. Science Translational Medicine. 9:eaal2774–eaal2774.http://www.ncbi.nlm.nih.gov/pubmed/29118260 http://www.pubmedcentral.nih.gov/articlerender.fcgi?artid=PMC5837001

Huang, X.Y., Z.L. Huang, J.H. Yang, Y.H. Xu, J.S. Sun, Q. Zheng, C. Wei, W. Song, and Z. Yuan. 2016. Pancreatic cancer cell-derived IGFBP-3 contributes to muscle wasting. J Exp Clin Cancer Res. 35:46.http://www.ncbi.nlm.nih.gov/pubmed/26975989

Keller, H.L., B. St Pierre Schneider, L.A. Eppihimer, and J.G. Cannon. 1999. Association of IGF-I and IGF-II with myofiber regeneration in vivo. Muscle Nerve. 22:347–354.http://www.ncbi.nlm.nih.gov/pubmed/10086895

Kim, D., B. Langmead, and S.L. Salzberg. 2015. HISAT: a fast spliced aligner with low memory requirements. Nat Methods. 12:357–360.https://www.ncbi.nlm.nih.gov/pubmed/25751142

Kirk, S.P., J.M. Oldham, F. Jeanplong, and J.J. Bass. 2003. Insulin-like growth factor-II delays early but enhances late regeneration of skeletal muscle. J Histochem Cytochem. 51:1611–1620.http://www.ncbi.nlm.nih.gov/pubmed/14623929

Kitamura, T., Y. Koshino, F. Shibata, T. Oki, H. Nakajima, T. Nosaka, and H. Kumagai. 2003. Retrovirus-mediated gene transfer and expression cloning: powerful tools in functional genomics. Exp Hematol. 31:1007–1014.http://www.ncbi.nlm.nih.gov/entrez/query.fcgi?cmd=Retrieve&db=PubMed&dopt=Citation&list_uids=14585362

Knight, J.D., and R. Kothary. 2011. The myogenic kinome: protein kinases critical to mammalian skeletal myogenesis. Skelet Muscle. 1:29.https://www.ncbi.nlm.nih.gov/pubmed/21902831

Kou, K., and P. Rotwein. 1993. Transcriptional activation of the insulin-like growth factor-II gene during myoblast differentiation. Mol Endocrinol. 7:291–302.http://www.ncbi.nlm.nih.gov/pubmed/8469241

Lefta, M., G. Wolff, and K.A. Esser. 2011. Circadian rhythms, the molecular clock, and skeletal muscle. Curr Top Dev Biol. 96:231–271.http://www.ncbi.nlm.nih.gov/pubmed/21621073

Levinovitz, A., E. Jennische, A. Oldfors, D. Edwall, and G. Norstedt. 1992. Activation of insulin-like growth factor II expression during skeletal muscle regeneration in the rat: correlation with myotube formation. Mol Endocrinol. 6:1227–1234.http://www.ncbi.nlm.nih.gov/pubmed/1406701

Liao, Y., G.K. Smyth, and W. Shi. 2014. featureCounts: an efficient general purpose program for assigning sequence reads to genomic features. Bioinformatics. 30:923–930.https://www.ncbi.nlm.nih.gov/pubmed/24227677

Lowe, M., J. Lage, E. Paatela, D. Munson, R. Hostager, C. Yuan, N. Katoku-Kikyo, M. Ruiz-Estevez, Y. Asakura, J. Staats, M. Qahar, M. Lohman, A. Asakura, and N. Kikyo. 2018. Cry2 is critical for circadian regulation of myogenic differentiation by Bclaf1-mediated mRNA stabilization of cyclin D1 and Tmem176b. Cell Rep. 22:2118–2132, PMCID: PMC29466738.http://www.ncbi.nlm.nih.gov/pubmed/29466738

Mayeuf-Louchart, A., B. Staels, and H. Duez. 2015. Skeletal muscle functions around the clock. Diabetes Obes Metab. 17 Suppl 1:39–46.http://www.ncbi.nlm.nih.gov/pubmed/26332967

McCarthy, J.J., J.L. Andrews, E.L. McDearmon, K.S. Campbell, B.K. Barber, B.H. Miller, J.R. Walker, J.B. Hogenesch, J.S. Takahashi, and K.A. Esser. 2007. Identification of the circadian transcriptome in adult mouse skeletal muscle. Physiol Genomics. 31:86–95.https://www.ncbi.nlm.nih.gov/pubmed/17550994

Mi, H., A. Muruganujan, and P.D. Thomas. 2013. PANTHER in 2013: modeling the evolution of gene function, and other gene attributes, in the context of phylogenetic trees. Nucleic Acids Res. 41:D377–386.https://www.ncbi.nlm.nih.gov/pubmed/23193289

Miller, B.H., E.L. McDearmon, S. Panda, K.R. Hayes, J. Zhang, J.L. Andrews, M.P. Antoch, J.R. Walker, K.A. Esser, J.B. Hogenesch, and J.S. Takahashi. 2007. Circadian and CLOCK-controlled regulation of the mouse transcriptome and cell proliferation. Proc Natl Acad Sci U S A. 104:3342–3347.https://www.ncbi.nlm.nih.gov/pubmed/17360649

Modi, H., C. Jacovetti, D. Tarussio, S. Metref, O.D. Madsen, F.P. Zhang, P. Rantakari, M. Poutanen, S. Nef, T. Gorman, R. Regazzi, and B. Thorens. 2015. Autocrine Action of IGF2 Regulates Adult beta-Cell Mass and Function. Diabetes. 64:4148–4157.http://www.ncbi.nlm.nih.gov/pubmed/26384384

Morita, S., T. Kojima, and T. Kitamura. 2000. Plat-E: an efficient and stable system for transient packaging of retroviruses. Gene Ther. 7:1063–1066.http://www.ncbi.nlm.nih.gov/entrez/query.fcgi?cmd=Retrieve&db=PubMed&dopt=Citation&list_uids=10871756

Motohashi, N., Y. Asakura, and A. Asakura. 2014. Isolation, culture, and transplantation of muscle satellite cells. J Vis Exp:doi: 10.3791/50846.http://www.ncbi.nlm.nih.gov/pubmed/24747722

Murphy, M.M., J.A. Lawson, S.J. Mathew, D.A. Hutcheson, and G. Kardon. 2011. Satellite cells, connective tissue fibroblasts and their interactions are crucial for muscle regeneration. Development. 138:3625–3637.http://www.ncbi.nlm.nih.gov/pubmed/21828091

Naumova, N., E.M. Smith, Y. Zhan, and J. Dekker. 2012. Analysis of long-range chromatin interactions using Chromosome Conformation Capture. Methods. 58:192–203.http://www.ncbi.nlm.nih.gov/pubmed/22903059

Paatela, E., D. Munson, and N. Kikyo. 2019. Circadian Regulation in Tissue Regeneration. Int J Mol Sci. 20.https://www.ncbi.nlm.nih.gov/pubmed/31071906

Pacheco-Bernal, I., F. Becerril-Perez, and L. Aguilar-Arnal. 2019. Circadian rhythms in the three-dimensional genome: implications of chromatin interactions for cyclic transcription. Clin Epigenetics. 11:79.http://www.ncbi.nlm.nih.gov/pubmed/31092281

Panda, S. 2016. Circadian physiology of metabolism. Science. 354:1008–1015.http://www.ncbi.nlm.nih.gov/pubmed/27885007

Papazyan, R., Y. Zhang, and M.A. Lazar. 2016. Genetic and epigenomic mechanisms of mammalian circadian transcription. Nat Struct Mol Biol. 23:1045–1052.http://www.ncbi.nlm.nih.gov/pubmed/27922611

Pizarro, A., K. Hayer, N.F. Lahens, and J.B. Hogenesch. 2013. CircaDB: a database of mammalian circadian gene expression profiles. Nucleic Acids Res. 41:D1009–1013.https://www.ncbi.nlm.nih.gov/pubmed/23180795

Rada-Iglesias, A., R. Bajpai, T. Swigut, S.A. Brugmann, R.A. Flynn, and J. Wysocka. 2011. A unique chromatin signature uncovers early developmental enhancers in humans. Nature. 470:279–283.http://www.ncbi.nlm.nih.gov/pubmed/21160473

Ren, H., P. Yin, and C. Duan. 2008. IGFBP-5 regulates muscle cell differentiation by binding to IGF-II and switching on the IGF-II auto-regulation loop. J Cell Biol. 182:979–991.http://www.ncbi.nlm.nih.gov/pubmed/18762576

Richards, J., B. Ko, S. All, K.Y. Cheng, R.S. Hoover, and M.L. Gumz. 2014. A role for the circadian clock protein Per1 in the regulation of the NaCl co-transporter (NCC) and the with-no-lysine kinase (WNK) cascade in mouse distal convoluted tubule cells. J Biol Chem. 289:11791–11806.http://www.ncbi.nlm.nih.gov/pubmed/24610784

Robinson, M.D., D.J. McCarthy, and G.K. Smyth. 2010. edgeR: a Bioconductor package for differential expression analysis of digital gene expression data. Bioinformatics. 26:139–140.https://www.ncbi.nlm.nih.gov/pubmed/19910308

Sasaki, H., P.A. Jones, J.R. Chaillet, A.C. Ferguson-Smith, S.C. Barton, W. Reik, and M.A. Surani. 1992. Parental imprinting: potentially active chromatin of the repressed maternal allele of the mouse insulin-like growth factor II (Igf2) gene. Genes Dev. 6:1843–1856.http://www.ncbi.nlm.nih.gov/pubmed/1383088

Segales, J., A.B. Islam, R. Kumar, Q.C. Liu, P. Sousa-Victor, F.J. Dilworth, E. Ballestar, E. Perdiguero, and P. Munoz-Canoves. 2016a. Chromatin-wide and transcriptome profiling integration uncovers p38alpha MAPK as a global regulator of skeletal muscle differentiation. Skelet Muscle. 6:9.http://www.ncbi.nlm.nih.gov/pubmed/26981231

Segales, J., E. Perdiguero, and P. Munoz-Canoves. 2016b. Regulation of Muscle Stem Cell Functions: A Focus on the p38 MAPK Signaling Pathway. Front Cell Dev Biol. 4:91.http://www.ncbi.nlm.nih.gov/pubmed/27626031

Shimizu-Motohashi, Y., Y. Asakura, N. Motohashi, N.R. Belur, M.G. Baumrucker, and A. Asakura. 2015. Pregnancy-induced amelioration of muscular dystrophy phenotype in mdx mice via muscle membrane stabilization effect of glucocorticoid. PLoS One. 10:e0120325.https://www.ncbi.nlm.nih.gov/pubmed/25775477

Siddle, K. 2011. Signalling by insulin and IGF receptors: supporting acts and new players. J Mol Endocrinol. 47:R1–10.http://www.ncbi.nlm.nih.gov/pubmed/21498522

Stow, L.R., J. Richards, K.Y. Cheng, I.J. Lynch, L.A. Jeffers, M.M. Greenlee, B.D. Cain, C.S. Wingo, and M.L. Gumz. 2012. The circadian protein period 1 contributes to blood pressure control and coordinately regulates renal sodium transport genes. Hypertension. 59:1151–1156.http://www.ncbi.nlm.nih.gov/pubmed/22526258

Takahashi, J.S. 2017a. Transcriptional architecture of the mammalian circadian clock. Nat Rev Genet. 18:164–179.https://www.ncbi.nlm.nih.gov/pubmed/27990019

Takahashi, J.S. 2017b. Transcriptional architecture of the mammalian circadian clock. Nature reviews. Genetics. 18:164–179.http://www.ncbi.nlm.nih.gov/pubmed/27990019 http://www.pubmedcentral.nih.gov/articlerender.fcgi?artid=PMC5501165

Taniguchi, C.M., B. Emanuelli, and C.R. Kahn. 2006. Critical nodes in signalling pathways: insights into insulin action. Nat Rev Mol Cell Biol. 7:85–96.http://www.ncbi.nlm.nih.gov/pubmed/16493415

Verma, M., Y. Asakura, B.S.R. Murakonda, T. Pengo, C. Latroche, B. Chazaud, L.K. McLoon, and A. Asakura. 2018. Muscle Satellite Cell Cross-Talk with a Vascular Niche Maintains Quiescence via VEGF and Notch Signaling. Cell Stem Cell. 23:530–543 e539.http://www.ncbi.nlm.nih.gov/pubmed/30290177

Wolff, C.A., and K.A. Esser. 2019. Exercise Timing and Circadian Rhythms. Curr Opin Physiol. 10:64–69.http://www.ncbi.nlm.nih.gov/pubmed/31938759

Yeung, J., J. Mermet, C. Jouffe, J. Marquis, A. Charpagne, F. Gachon, and F. Naef. 2018. Transcription factor activity rhythms and tissue-specific chromatin interactions explain circadian gene expression across organs. Genome Res. 28:182–191.http://www.ncbi.nlm.nih.gov/pubmed/29254942

Yoon, Y.S., S. Jeong, Q. Rong, K.Y. Park, J.H. Chung, and K. Pfeifer. 2007. Analysis of the H19ICR insulator. Mol Cell Biol. 27:3499–3510.http://www.ncbi.nlm.nih.gov/pubmed/17339341

Yoshiko, Y., K. Hirao, and N. Maeda. 2002. Differentiation in C(2)C(12) myoblasts depends on the expression of endogenous IGFs and not serum depletion. Am J Physiol Cell Physiol. 283:C1278–1286.http://www.ncbi.nlm.nih.gov/pubmed/12225990

Zhang, Y., T. Liu, C.A. Meyer, J. Eeckhoute, D.S. Johnson, B.E. Bernstein, C. Nusbaum, R.M. Myers, M. Brown, W. Li, and X.S. Liu. 2008. Model-based analysis of ChIP-Seq (MACS). Genome Biol. 9:R137.http://www.ncbi.nlm.nih.gov/pubmed/18798982

Zheng, B., U. Albrecht, K. Kaasik, M. Sage, W. Lu, S. Vaishnav, Q. Li, Z.S. Sun, G. Eichele, A. Bradley, and C.C. Lee. 2001. Nonredundant roles of the mPer1 and mPer2 genes in the mammalian circadian clock. Cell. 105:683–694.https://www.ncbi.nlm.nih.gov/pubmed/11389837

Zheng, B., D.W. Larkin, U. Albrecht, Z.S. Sun, M. Sage, G. Eichele, C.C. Lee, and A. Bradley. 1999. The mPer2 gene encodes a functional component of the mammalian circadian clock. Nature. 400:169–173.http://www.ncbi.nlm.nih.gov/pubmed/10408444

